# Integrated Organ Immunity: Antigen-specific CD4-T cell-derived IFN-γ induced by BCG imprints prolonged lung innate resistance against respiratory viruses

**DOI:** 10.1101/2023.07.31.551354

**Authors:** Audrey Lee, Katharine Floyd, Shengyang Wu, Zhuoqing Fang, Tze Kai Tan, Chunfeng Li, Harold Hui, David Scoville, Alistaire D. Ruggiero, Yan Liang, Anna Pavenko, Victor Lujan, Garry P. Nolan, Prabhu Arunachalam, Mehul Suthar, Bali Pulendran

## Abstract

Bacille Calmette-Guérin (BCG) vaccination can confer non-specific protection against heterologous pathogens. However, the underlying mechanisms remain mysterious. Here, we show that mice immunized intravenously with BCG exhibited reduced weight loss and/or improved viral clearance when challenged with SARS-CoV-2 and influenza. Protection was first evident between 14 - 21 days post vaccination, and lasted for at least 42 days. Remarkably, BCG induced a biphasic innate response in the lung, initially at day 1 and a subsequent prolonged phase starting at ∼15 days post vaccination, and robust antigen-specific Th1 responses. MyD88-dependent TLR signaling was essential for the induction of the innate and Th1 responses, and protection against SARS-CoV-2. Depletion of CD4^+^ T cells or IFN-γ activity prior to infection obliterated innate activation and protection. Single cell and spatial transcriptomics revealed CD4-dependent expression of interferon-stimulated genes (ISGs) in myeloid, type II alveolar and lung epithelial cells. Thus, BCG elicits “integrated organ immunity” where CD4+ T cells act on local myeloid and epithelial cells to imprint prolonged antiviral innate resistance.

## INTRODUCTION

Vaccines provide protection against pathogens primarily through the generation of antigen-specific antibodies or cytotoxic T cell responses^1^. However, epidemiological studies conducted over the years have suggested that certain live attenuated vaccines, such as the oral polio or Bacille Calmette-Guérin (BCG) vaccine, might also confer non-specific protection against unrelated infections, as indicated by a reduction in infant mortality following vaccination^2–5^. Remarkably, the recent COVID-19 pandemic has drawn attention to the BCG vaccine, with emerging studies in human controlled trials and murine models highlighting its effectiveness in providing non-specific antiviral effects against the SARS-CoV-2 virus^6–9^. However, the efficacy of BCG vaccine in the non-specific protection against SARS-CoV-2 remains contentious, as conflicting findings have been presented in other studies^10–12^.

The BCG vaccine, which contains live attenuated *Mycobacterium bovis*, is widely used globally in the prevention against tuberculosis (TB), particularly in areas where TB is prevalent. Despite its extensive use, the precise mechanism by which the BCG vaccine confers protection against TB, let alone diverse pathogens, remains poorly understood. While CD4^+^ T cells, including Th1 and Th17 cells, have been recognized as important immune mediators in TB protection^13–15^, other studies have not consistently identified similar immune correlates^16,17^. Moreover, several studies have highlighted the synergistic involvement of CD8^+^ T cells and T cell-independent mechanisms, such as activated alveolar macrophages, in contributing to optimal immunity against TB^18–20^.

The emerging concept of trained immunity, whereby innate immune cells retain long-lasting memory in the form of epigenetic or metabolic reprogramming^21,22^, has been observed following BCG vaccination^23,24^. In fact, a recent study revealed that following intravenous BCG vaccination in mice, bone marrow stem and progenitor cells undergo long-term reprogramming, also known as innate training, and play a role in T cell-independent protection against TB^25^. Training of non-specific innate cells has thus been proposed to be responsible for the non-specific effects of the BCG vaccine. In line with this notion, recent studies have suggested that enhanced cytokine responses produced by trained myeloid cells following a secondary stimulation could be a key mediator in conferring protection against heterologous pathogens^26,27^.

While there have been mounting studies addressing the non-specific effects of BCG in the context of trained immunity, there remains a need for a comprehensive evaluation of BCG-induced immune responses and their direct impact on non-specific antiviral activity in vivo. A deeper understanding of the underlying mechanisms of heterologous protection is crucial for the rational design and development of universal vaccines aimed at preventing a broad range of infections. In this study, we delineated the mechanism underlying the non-specific antiviral effects exerted by the BCG vaccine against SARS-CoV-2 virus, and reveal a pivotal role for BCG-specific CD4^+^ T cells that produce IFN-γ in imprinting a persistent antiviral innate program in the lung, mediating heterologous viral protection.

## RESULTS

### BCG vaccination protects mice against challenge with SARS-CoV-2 and influenza PR8

To investigate the non-specific effects of BCG vaccine on protection against heterologous viruses, mice were administered with BCG intravenously (BCG IV) and challenged with SARS-CoV-2 B.1.351 virus at day 7, 14, 21, 28 or 42 post-vaccination. BCG had previously been shown to elicit enhanced immune responses and protection in non-human primate and murine models when administered intravenously^7,28,29^. We found that vaccinated mice were protected from weight loss when challenged at day 21, 28, 42 post-vaccination, compared to unvaccinated control mice **(Fig. 1a)**. Surprisingly however, the reduction in weight loss was not observed at days 7 or 14 post-vaccinated mice. In addition, mice vaccinated at days 21, 28 and 42 post-prior to challenge with SARS-CoV-2 exhibited ∼41-, ∼435-, and ∼92-fold reduction in lung viral load respectively, as measured by the expression of SARS-CoV-2 RNA-dependent RNA polymerase gene at day 3 post-challenge **(Fig. 1b)**. Notably, the reduction in viral load was mainly observed in the lungs and not the nasal turbinates **(Fig. 1b)**, suggesting that protective effects were largely confined to the lungs and not the upper respiratory tract. We then performed hematoxylin and eosin (H&E) of lung tissues to assess the level of pathology following challenge. We found that lung inflammation was significantly reduced in mice challenged at 21 days post-vaccination, although this was not observed in day 28- or 42-vaccinated mice **(Fig 1c)**. Immunohistochemical staining of the lungs post-challenge further revealed a lower expression of SARS-CoV-2 nucleocapsid protein (NP) that localized to alveolar type I and II pneumocytes, alveolar macrophages, and diffuse bronchiolar epithelial cells in mice challenged at day 21, 28 and 42 post-vaccination **(Fig. 1c and Supplementary Fig. 1a)**, suggesting that these cell types largely contributed to the reduction in viral load observed in BCG-vaccinated mice. Together, these findings suggest that BCG provided a temporal window of protection against SARS-CoV-2, beginning prior to 21 days following vaccination and lasting at least as long as 42 days.

**Figure 1.**
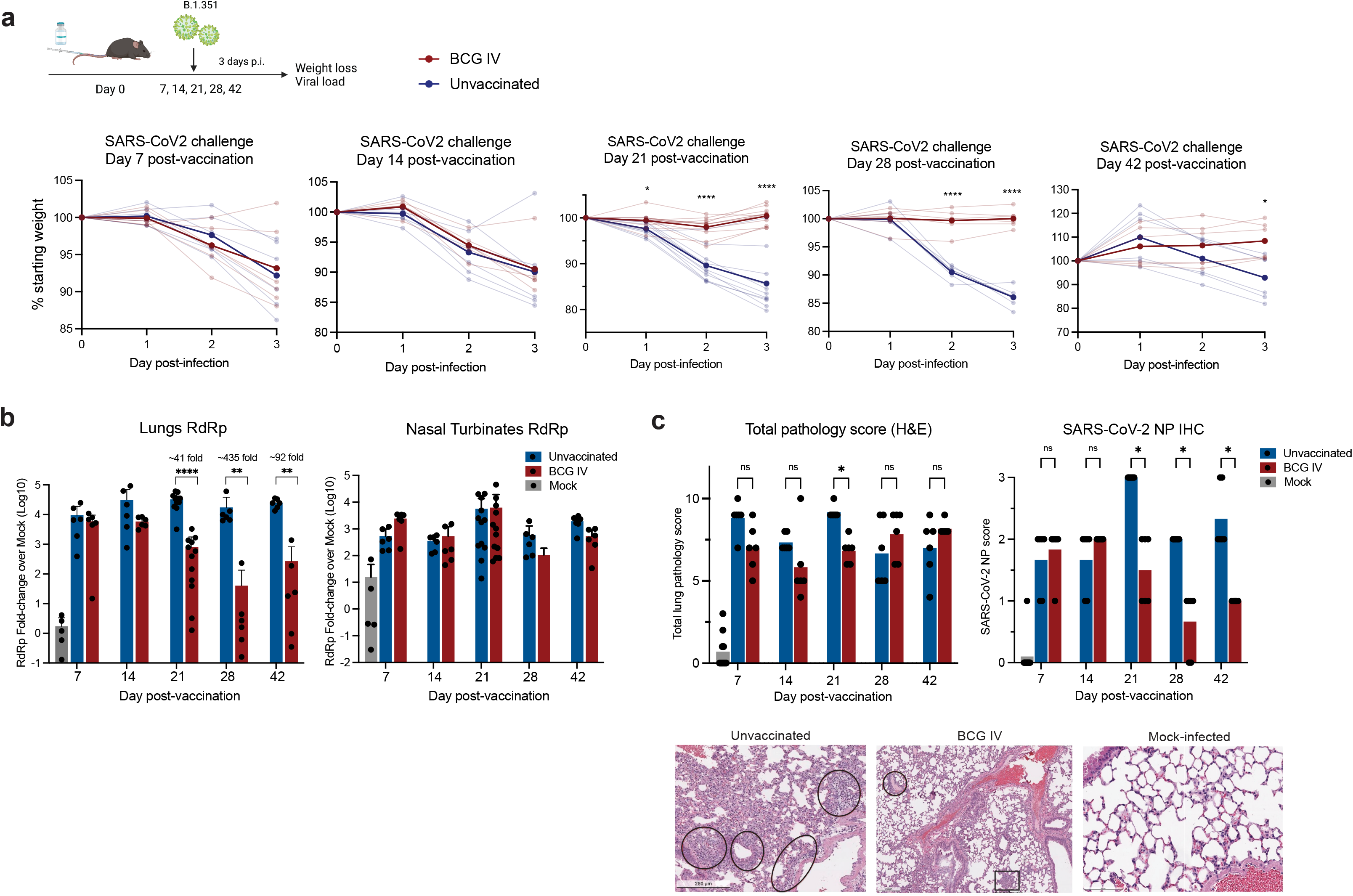
BCG vaccination protects mice against challenge with SARS-CoV-2. **a**, Experimental outline and weight loss of BCG IV vaccinated and unimmunized control mice following SARS-CoV-2 challenge at day 7, 14, 21, 28, 42 post-vaccination. Mean % starting weight and % starting weight of individual mice are shown here. **b,** Log10 fold-change of SARS-CoV-2 RNA-dependent RNA polymerase (RdRp) gene in lungs and nasal turbinates of vaccinated or unvaccinated mice over mock-infected mice, as measured by RT-PCR. **c,** Pathology score and SARS-CoV-2 nucleocapsid protein (NP) score of lung tissue sections measured at D3 post-challenge. Bottom, Representative H&E stained lung sections from unvaccinated, BCG IV, mock-infected mice with circles and rectangle representing peribronchiolar, perivascular inflammation and alveolar inflammation, respectively. In panel **a and b**, data combined from two independent experiments at D21 (n=12) and representative of one independent experiment at D7, 14, 28, 42 (n=6). In **c**, data representative of two independent experiments. Statistical analysis was performed by Two-way ANOVA with Sidak’s multiple comparisons test **(a)**, multiple Mann-Whitney with Holm-Sidak multiple comparisons test **(b and c)**. *, P < 0.05; **, P < 0.01; ****, P < 0.0001.

To assess if viral protection was limited to intravenous administration of the vaccine, we vaccinated mice with BCG via the intranasal route prior to SARS-CoV-2 B.1.351 challenge. Interestingly, we found that mice given intranasal BCG similarly showed a reduction in weight loss and lung viral load, although not statistically significant **(Supplementary Fig. 1b)**. Additionally, to test the extent of non-specific protection against other pathogens, we vaccinated mice via intravenous (IV), intranasal (IN), intramuscular (IM), and subcutaneous (SC) routes and infected them with a lethal dose of the Influenza A H1N1/PR8 virus (PR8) at day 21 post-vaccination. We measured their weight loss and survival up till 14 days of challenge. Interestingly, mice vaccinated via IV, IM, IN routes exhibited reduced weight loss (percentage of initial body weight at day 6 post-challenge, IV=93.3%; IN=94.1%; IM=88.6%; unvaccinated=81.7%) and improved survival (IV= ∼18.2%; IN=∼75%; IM=∼17.6%; unvaccinated=0%), as compared to unvaccinated control mice **(Supplementary Fig. 1c)**. However, mice vaccinated via the subcutaneous route had no effect on survival rate or weight loss **(Supplementary Fig. 1c)**. These results demonstrated that protection conferred by BCG was dependent on the route of vaccination.

### BCG vaccination induced a biphasic pattern of innate activation

To understand the mechanism underlying the non-specific effects of BCG, we assessed the kinetics of cytokine and cellular immune responses in mice following BCG IV vaccination. Analysis of serum cytokines in BCG-vaccinated mice revealed a biphasic pattern in the kinetics of production of systemic inflammatory mediators, with an early peak of innate cytokines and chemokines (i.e. TNF-α, MCP3, CCL2, GCSF, GM-CSF) at 6 hours post-vaccination and a late peak (i.e. MCP3, IFN-γ, TNF-α, RANTES) emerging around 21 days after vaccination **(Fig. 2a and Supplementary Fig. 2a)**. Furthermore, the frequency of CD11b^+^ DCs and activation of Ly6C^+^ monocytes, CD11b^+^ and CD103^+^ DCs in the lungs appeared to follow a similar biphasic pattern **(Fig. 2b)**. Specifically, the frequency of CD11b^+^ DCs was significantly increased at day 1 and around day 15 post-vaccination. Likewise, in Ly6C^+^ monocytes, CD11b^+^ and CD103^+^ DCs, CD86 activation marker was upregulated at day 1 and day 15 to 21 post-vaccination, and in the case of Ly6C^+^ monocytes and CD11b DCs, maintained at least as long as day 42 **(Fig. 2b)**. Interestingly, lung interstitial macrophages (F4/80^+^ macrophages) increased in frequency early at day 1, whereas the frequency of alveolar macrophages decreased beginning day 1, although their activation was enhanced till day 42 post-vaccination **(Fig. 2b and Supplementary Fig. 2b)**.

**Figure 2.**
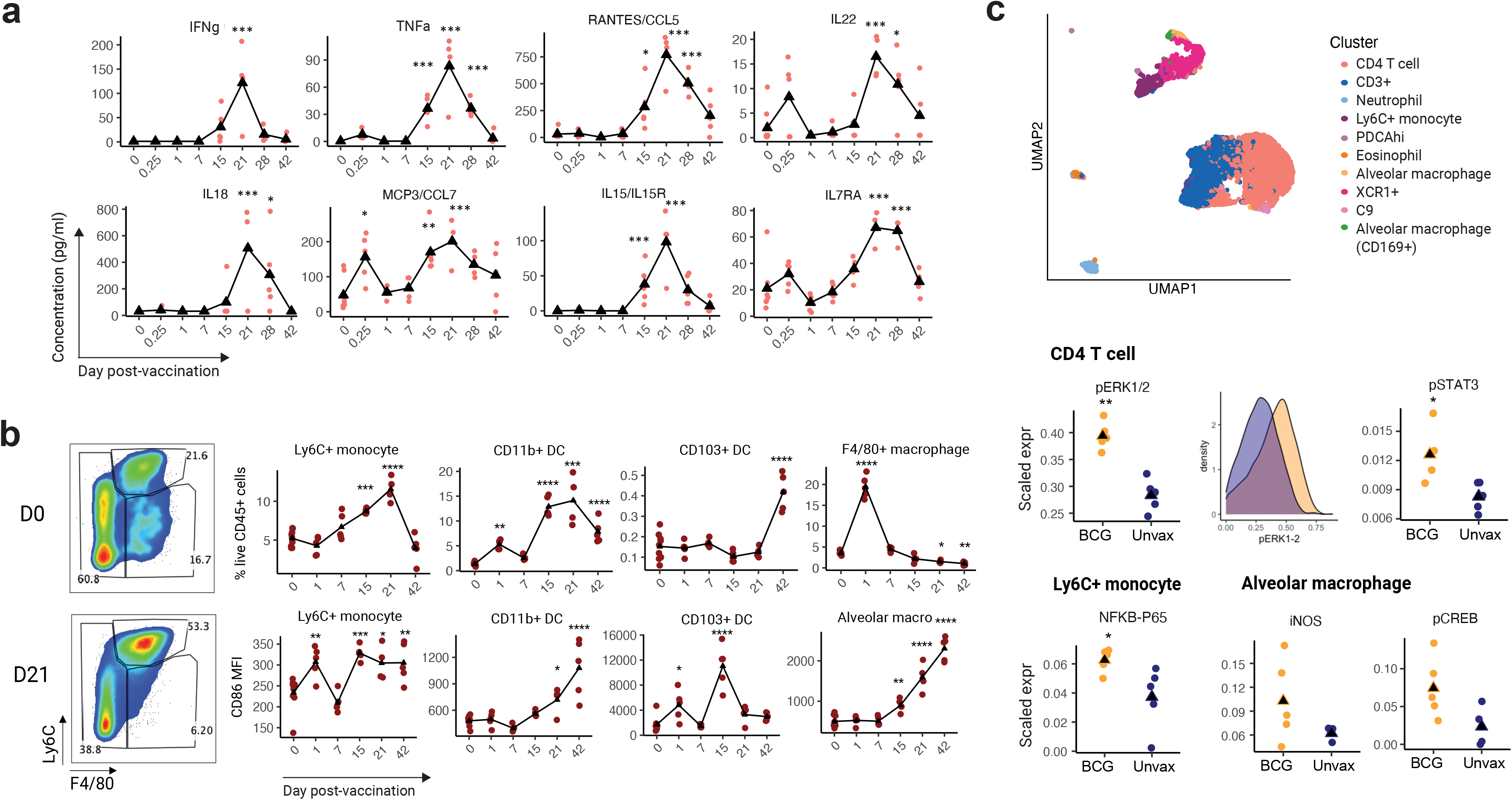
BCG vaccination induced a biphasic pattern of innate activation. **a**, Serum cytokine measured by Luminex assay at various timepoints post-BCG IV vaccination (n=4-5). Data representative of two independent experiments. **b**, Frequency (% live CD45+ cells) and activation indicated by CD86 median fluorescence intensity (MFI) of innate cell populations in the lungs post-BCG IV vaccination (n=4-5). Data representative of two independent experiments at day 21. **c**, top, CyTOF analysis of lung immune cells at day 21 post-vaccination. Uniform Manifold Approximation and Projection (UMAP) of 17,638 total live cells obtained in the lungs of unvaccinated (unvax) and BCG IV vaccinated mice. **bottom,** Normalized and scaled mean values of intracellular markers found in CD4 T cells, Ly6C+ monocytes, and alveolar macrophages (AM) in individual mice, visualized as dot plots and distribution plot (pERK1-2 in CD4+ T cells). Wilcoxon rank-sum test. Data from one representative experiment. In panel **a** and **b**, One-way ANOVA with Tukey’s multiple comparisons test **(a)** and Dunnett’s multiple comparison test for each timepoint compared to day 0 **(b).** *, P < 0.05; **, P < 0.01; ***, P < 0.005; ****, P < 0.0001.

Similarly, we found an increased proportion of macrophages in the spleen at day 1 and increased frequencies and activation of Ly6C^+^ monocytes, CD11b^+^ and CD8+ DCs beginning 15 days post-vaccination **(Supplementary Fig. 2c)**. Reprogramming and activation of bone marrow stem cells have previously been associated with sustained training of myeloid cells^25,30^. To identify activity in the bone marrow, we analyzed the kinetics of bone marrow stem or progenitor cells. We found that at 15 to 42 days post-vaccination, there were enhanced frequencies of Lin^−^Sca-1^+^c-Kit^+^ (LSK) cells, myeloid-biased multipotent progenitors (MPP3 and MPP2), and short-term haematopoietic stem cells (ST-HSCs), suggesting active bone marrow proliferation at these timepoints **(Supplementary Fig. 2d)**.

To obtain further insights into the nature of the innate response at day 21 post-vaccination, we used mass cytometry (CyTOF) with a panel of key surface and intracellular signaling proteins **(Supplementary Fig. 3).** We identified upregulation of the NF-kB p65 subunit in Ly6C^+^ monocyte population and increased expression of phosphorylated ERK1/2 and phosphorylated STAT3, which are signaling proteins important for T cell activation and IL-10 production^31^, in CD4 T cells in the lung **(Fig. 2c)**. A trend for increased expression of inducible nitric oxide (iNOS) and phosphorylated CREB in alveolar macrophages was also observed in the lungs **(Fig. 2c)**. Similarly, upregulation of phosphorylated IRF3, STAT1, ERK1/2, and NF-kB p65 and STAT5, was observed in splenic CD4^+^ T cells and Ly6C^+^ monocytes, respectively **(Supplementary Fig. 3)**. Together, these findings demonstrate that BCG vaccination induces systemic immune activation and pro-inflammatory responses at 21 to 42 days.

### BCG vaccination stimulates an antigen-specific Th1 response

We next profiled the BCG-specific T cell responses in the lung and spleen following vaccination. At day 21 post-vaccination, there was a significant expansion of BCG-specific CD4^+^ T cells secreting IFN-γ, TNF-α, and/or IL-2, but these cells were not present at day 7 post-vaccination **(Fig. 3a -c)**. Similarly, CD4^+^ T cells induced in the lungs and spleen were predominantly of the Th1 subset that are either single producers of IFN-γ or double producers of IFN-γ and TNF-α **(Fig. 3b, d)**. Collectively, these findings suggest that intravenous BCG induced a systemic Th1 response in both the lung and spleen.

**Figure 3.**
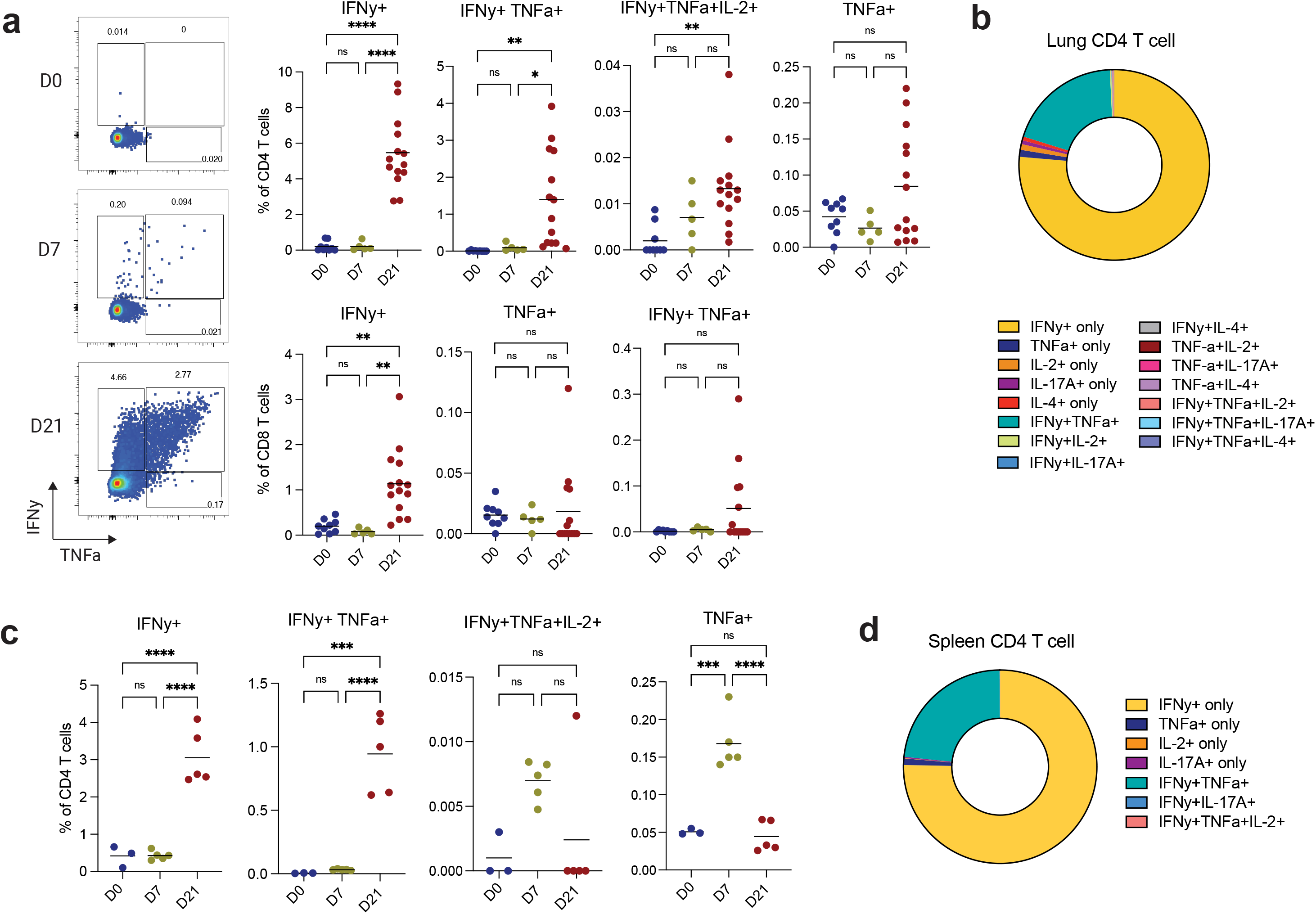
BCG vaccination induced a robust Th1 profile at 21 days post-vaccination. **a**, Lung CD4 and CD8 T cell responses at baseline, day 7, 21 post-BCG IV vaccination. Left, representative flow cytometry plot of CD4 T cells expressing IFN-g or TNF-a. D0 and D21 data combined from 2 independent experiments (D0, n=9; D21, n=14). Day 7 data from 1 independent experiment (n=5). **b,** Lung CD4 T cell proportion of T cell subsets. **c,** Spleen CD4 T cell responses at baseline, day 7, 21 post-BCG IV vaccination. **d,** Spleen CD4 T cell proportion of T cell subsets (D0, n=3; D7 and D21, n=5). Statistical analysis was performed by One-way ANOVA with Tukey multiple comparisons test. *, P < 0.05; **, P < 0.01; ***, P < 0.005; ****, P < 0.0001.

### BCG vaccination protects against SARS-CoV-2 via MyD88 signaling

BCG is a bacterium that contains different pathogen-associated molecular patterns (PAMPS) and is known to activate multiple innate sensing pathways, such as TLR-MyD88 and inflammasome pathways^32^ **(Fig. 4a)**. To investigate the nature of the innate mechanisms contributing to non-specific protection against SARS-CoV-2 B.1.351, we challenged various knock-out mice deficient in MyD88, the inflammasome component ASC, and STING pathways (*Myd88^−/−−^*, *Asc^−/−^*, *Sting^−/−^* mice respectively). We found that the reduced weight loss observed in BCG-vaccinated wild-type (WT) mice upon SARS-CoV-2 B.1.351 challenge was completely abrogated in *Myd88^−/−^* mice, whereas *Asc^−/−^* and *Sting^−/−^* mice exhibited similar weight loss as vaccinated WT control **(Fig. 4b)**. The abrogation of protection against SARS-Cov-2 seen in vaccinated *Myd88^−/−^* mice was reflected by a concomitant lack of reduction in the lung viral loads in such mice **(Fig. 4c).** *Myd88^−/−^* abrogated the reduction of viral load in the lungs, whereas *Asc^-/^*, and *Sting^−/−^* vaccinated mice showed a reduction in SARS-CoV-2 B.1.351 viral load upon challenge, albeit at a lower fold-change (*Asc^−/−^*: ∼7.8-fold; *Sting^−/−^*: ∼19-fold) compared to WT control (∼41-fold) **(Fig. 4c)**. Consistently, there was no difference in the lung pathology and SARS-CoV-2 nucleocapsid protein level at day 3 post-challenge in vaccinated compared to unvaccinated groups in *Myd88^−/−^* mice **(Fig. 4d)**. In contrast, *Asc^−/−^* and *Sting^−/−^* mice administered the BCG vaccine exhibited a milder lung pathology and SARS-CoV-2 nucleocapsid protein level **(Fig. 4d)**. These results demonstrated that the protective effect of BCG against SARS-CoV-2 was largely dependent on *Myd88*-signaling, but not on *Asc*- or *Sting*-dependent pathways.

**Figure 4.**
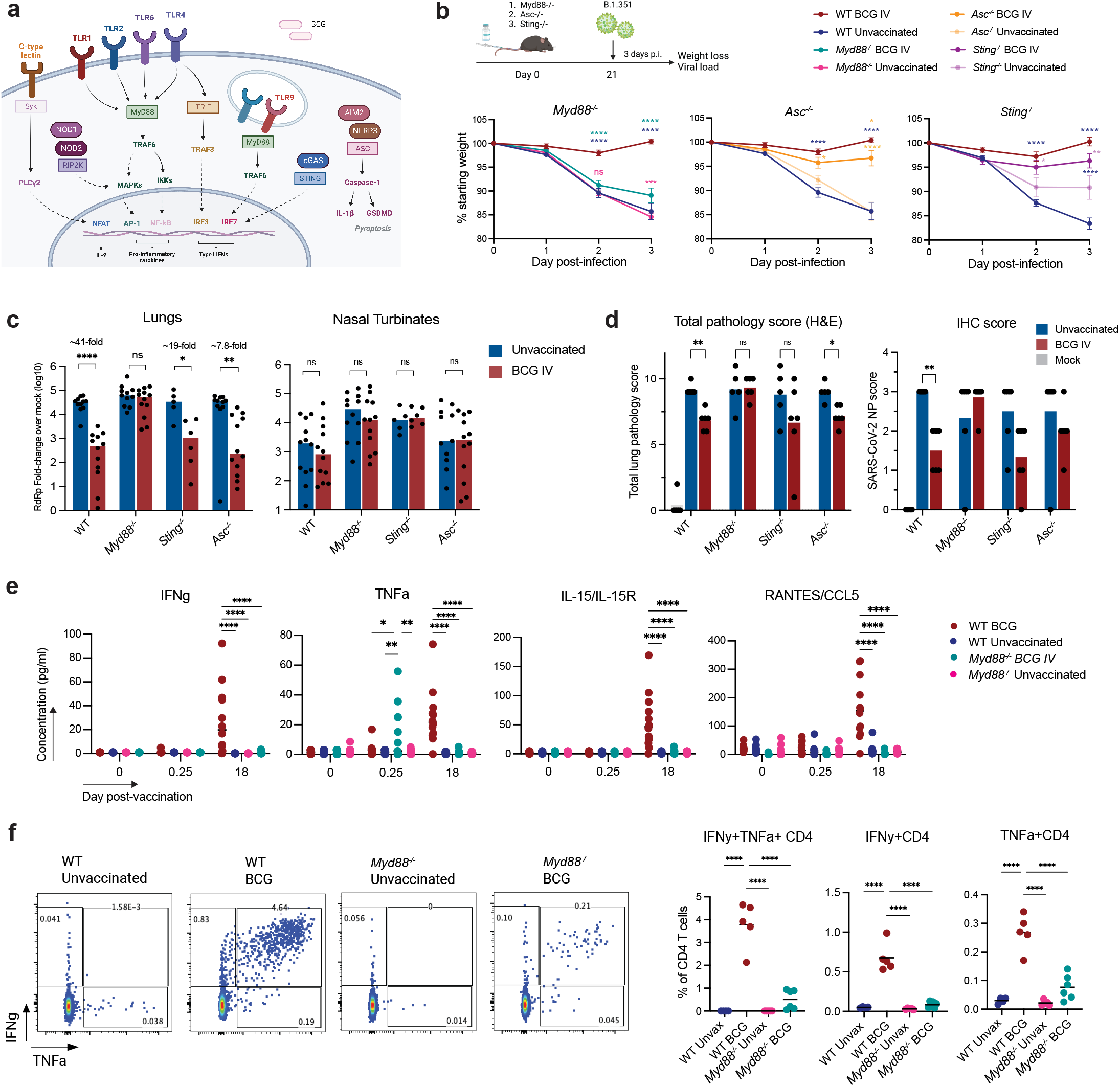
Protection against SARS-CoV-2 B.1.351 conferred by BCG vaccination is dependent on MyD88 signaling. **a**, Diagram depicting the innate signaling pathways induced by BCG bacterium. **b,** Experimental outline and weight loss of vaccinated and unvaccinated knock-out mice following SARS-CoV-2 challenge. Data shown is mean±SEM. **c,** SARS-CoV-2 viral load in lungs and nasal turbinates at D3 post-challenge expressed as fold-change over mock-infected mice. In panel **b** and **c**, Wild-type (WT), *Myd88^−/−^* and *Asc^−/−^* data combined from two independent experiments (n=11-12). *Sting^−/−^* data representative of one independent experiment (n=5-6). **d,** Pathology score and SARS-CoV-2 nucleocapsid protein (NP) score of lung tissue sections measured at D3 post-challenge in knock-out models (n=6). **e,** Serum cytokine responses in BCG IV and unvaccinated WT and *Myd88^−/−^* mice pre-challenge. Data combined from two independent experiments (n=12-13). **f,** T cell responses in BCG IV and unvaccinated *Myd88^−/−^* compared to WT mice. Data representative of two independent experiments (n=5-6). Statistical analysis was performed by Two-way ANOVA with Tukey’s multiple comparison test (panel **b, e, f**). Multiple Mann-Whitney with Holm-Sidak multiple comparisons test (panel **c, d**). *, P < 0.05; **, P < 0.01; ****, P < 0.0001.

To better understand the differential protection outcome in mice lacking MyD88 signaling, we analyzed the immune responses to BCG in *Myd88^−/−^* and WT mice. We found that pro-inflammatory cytokines, such as IFN-γ, TNF-α, IL-15, CCL5 and CXCL10, were barely detectable in the serum at day 18 post-vaccination in *Myd88^−/−^*, compared to vaccinated WT mice **(Fig. 4e and Supplementary Fig. 4a)**. However, *Myd88^−/−^* mice were not entirely unresponsive to the BCG vaccine, as some chemokines, such as CXCL10 and CCL2, remained detected at 6 hours post-vaccination, although at a level lower than vaccinated WT mice **(Supplementary Fig. 4a)**. These differences in serum level of pro-inflammatory cytokines between vaccinated *Myd88^−/−^* and WT mice were also observed 3 days after viral challenge **(Supplementary Fig. 4b)**. Conversely, *Asc^−/−^*, and *Sting^−/−^* mice resulted in a modest decrease of IFN-γ and TNF-α production that is not as profound as *Myd88^−/−^* mice, when compared to WT mice in response to BCG **(Supplementary Fig. 4c, d)**.

Lastly, we compared the T cell responses to BCG vaccination in *Myd88^−/−^* and WT mice. We found that the significant increase in the frequency of IFN-γ^+^, TNF-α^+^ or IL-2^+^ CD4^+^ T cells in the lungs of WT mice at day 21 post-vaccination was profoundly reduced in *Myd88^−/−^* mice **(Fig. 4f)**. Collectively, these findings demonstrate that the innate and antigen-specific T cell response and heterologous protection conferred by BCG was largely dependent on the MyD88 signaling pathway.

### CD4^+^ T cells and IFN-γ production play a critical role in the heterologous protection mediated by BCG vaccination

Given that *Myd88^−/−^* mice had severe impairments in the generation of BCG-specific CD4^+^ T cell response, serum cytokines, and heterologous protection against SARS-CoV-2 **(Fig. 4)**, and given the synchronized kinetics of the CD4^+^ T cell response and second phase of the innate response, we postulated that cytokine feedback from the CD4^+^ T cells may stimulate the second phase of innate activation. To test this, we depleted either CD4, CD8, or both CD4 and CD8 T cells (CD4/CD8) in vaccinated mice by injecting anti-CD4 or anti-CD8 antibodies every 2 consecutive days, beginning 5 days prior to viral challenge **(Fig. 5a)**. We found that mice depleted of CD4^+^ T cells were unable to reduce viral load in the lungs and mice depleted of both CD4^+^ and CD8^+^ T cells had only a modest reduction in viral load as compared to vaccinated WT mice, whereas mice depleted of only CD8^+^ T cells exhibited significant reduction in lung viral load **(Fig. 5a)**. The lack of protection in mice depleted of CD4^+^ T cells were further demonstrated in their weight loss that is comparable to unvaccinated control following viral challenge **(Fig. 5b)**. To exclude the possibility that the observed phenotype in CD4-depleted vaccinated mice was a consequence of SARS-CoV-2 specific CD4^+^ T cells orchestrating adaptive immunity against SARS-CoV-2, we tested the effects of antibody depletion on viral challenge outcome in unvaccinated control mice and observed that CD4 and/or CD8 depletion did not affect the challenge outcome in unvaccinated mice **(Supplementary Fig. 5)**. This is in line with a previous study indicating that SARS-CoV-2 viral load early in infection was unaffected by the absence of T cells^33^.

**Figure 5.**
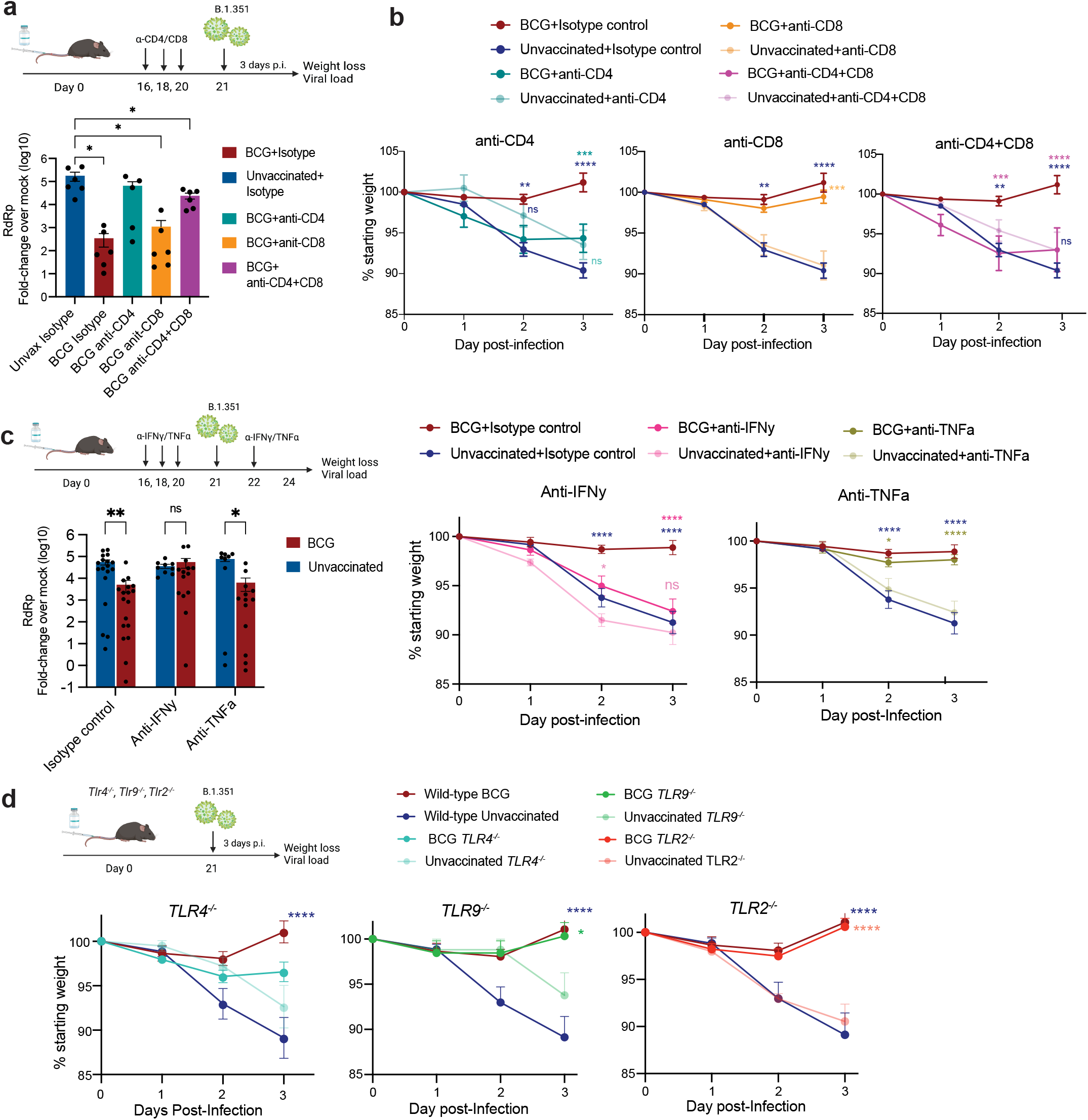
CD4 T cells and IFN-γ production play a critical role in the heterologous protection mediated by BCG vaccination. **a**, Experimental outline and lung SARS-CoV-2 viral load following T cell depletion and challenge. Data representative of two independent experiments (n=6). **b,** Weight loss of BCG IV vaccinated and unimmunized control mice following T cell depletion and SARS-CoV-2 challenge. Data combined from two independent experiments (n=6-12). **c,** Lung viral load and weight loss from SARS-CoV-2 challenge following cytokine depletion. Data combined from two independent experiments (n=9-18). **d,** Weight loss following SARS-CoV-2 challenge in *TLR4^−/−^, TLR9^−/−^, TLR2^−/−^* mice. Weight loss shown here are plotted as mean±SEM. Data combined from two independent experiments. Statistical analysis was performed by Two-way ANOVA with Tukey’s multiple comparison test (panel **b, c, d** weight loss). Multiple Mann-Whitney with Holm-Sidak multiple comparisons test (panel **a**). *, P < 0.05; **, P < 0.01; ****, P < 0.0001.

As most BCG-specific CD4^+^ T cells induced by vaccination were Th1 cells that produced IFN-γ and TNF-α, we determined if the lack of protection observed following CD4^+^ T cell depletion could be a result of diminished IFN-γ or TNF-a concentration *in vivo*. To achieve this, we administered anti-IFN-γ or anti-TNF-α antibodies every 2 consecutive days, beginning 5 days prior to viral challenge and 1 day post-challenge **(Fig. 5c)**. Depletion of IFN-γ abrogated the reduction in viral load and weight loss conferred by BCG vaccination **(Fig. 5c)**. However, TNF-α depletion had no effect on both weight loss and viral load **(Fig. 5c)**. These results demonstrate that CD4^+^ T cells and IFN-γ stimulated by BCG are critical mediators of viral protection.

We next determined the target receptor along the TLR-MyD88 signaling pathway that is crucial for inducing BCG-specific immune responses and heterologous viral protection. Since bacteria express ligands for TLRs 4, 2 and 9, we tested the SARS-CoV-2 B.1.351 challenge outcome on TLR4-, TLR9-, and TLR2-knockout mouse models (*Tlr4^−/−^*, *Tlr9^−/−^*, *Tlr2^−/−^*). Specifically, vaccinated *Tlr4^−/−^* mice did not exhibit improved weight loss following SARS-CoV-2 B.1.531 challenge, whereas vaccinated *Tlr9^−/−^* and *Tlr2^−/−^* displayed a similar reduction in weight loss to that of vaccinated WT mice **(Fig. 5d)**. These results suggest a mechanism whereby BCG activated the TLR4-Myd88 signaling pathway and induced Th1 responses that play a major role in mediating non-specific protection.

### Th1 cells and IFN-γ drive the activation of myeloid cells and lung epithelial cells

To further explore changes in the immune landscape within the lungs following BCG vaccination, we performed single cell RNA-sequencing (scRNA-seq) of whole lungs at day 0 and 21 post-vaccination. We obtained a total of 25,622 cells in the lungs and annotated distinct clusters of immune cells (myeloid, alveolar macrophage, NK and T cells) and non-immune cells consisting of lung epithelial (alveolar type I and II pneumocytes, club cells, pulmonary neuroendocrine cells; PNEC), endothelial and mesenchymal cells (alveolar fibroblast, airway smooth muscle cells; airway SMC, mesothelial) **(Fig. 6a and Supplementary Fig. 6a)**.

**Figure 6.**
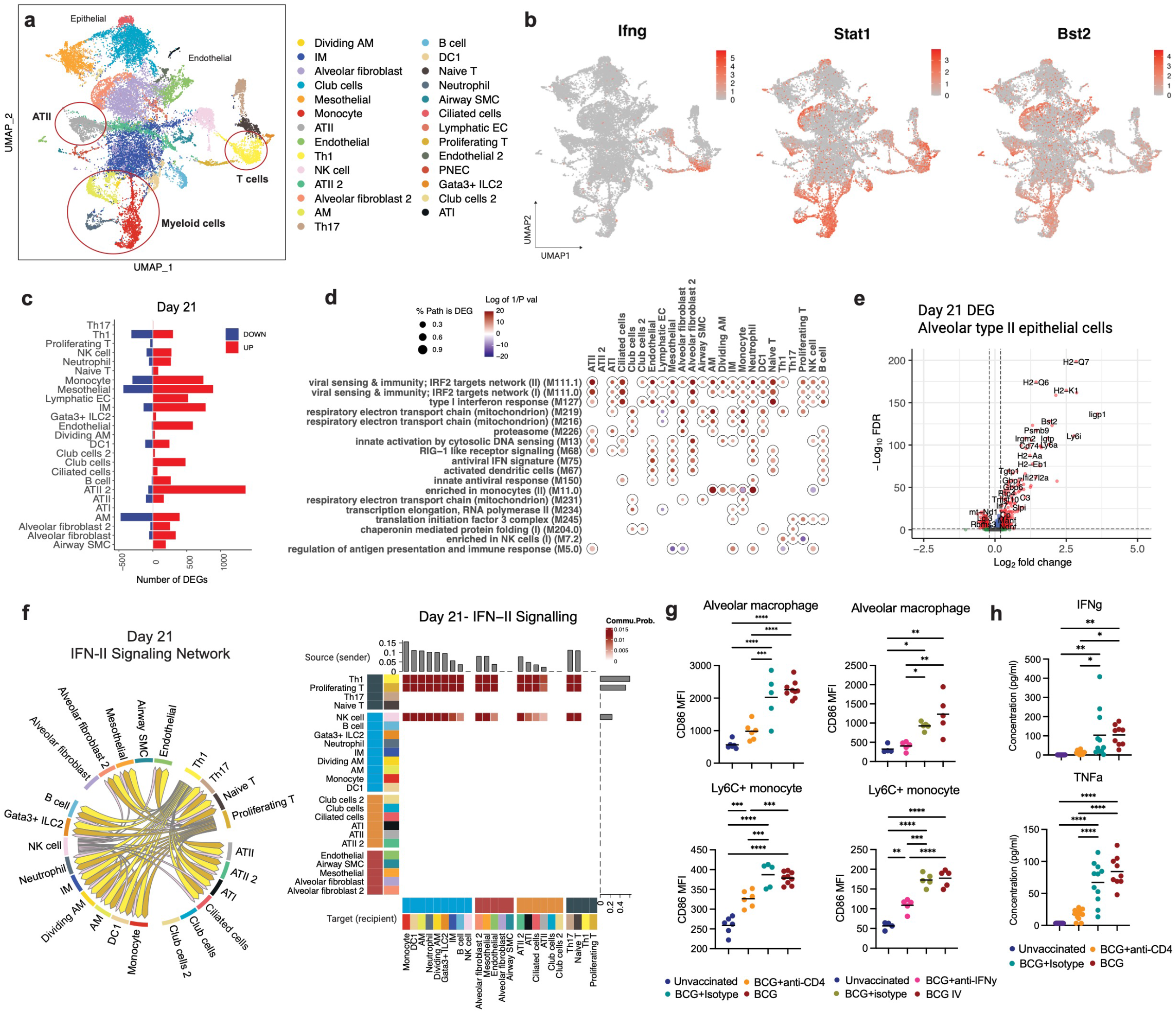
Th1 cells and IFN-γ drive the activation of myeloid cells and lung epithelial cells. **a**, UMAP projection of 25,622 total lung cells clustered from scRNA-seq of day 0 and 21 BCG IV vaccinated mice. Alveolar type II pneumocytes (ATII), myeloid cells, and CD4 T cells are highlighted in red circles. **b,** UMAP projection colored by scaled expression of markers. **c,** Differentially expressed genes (DEGs) at day 21 versus day 0 (absolute log2FC > 0.25, FDR < 0.05; differential analysis performed by Wilcoxon rank-sum test with Benjamini Hochberg (BH) correction). **d,** Blood transcriptional modules (BTMs) enrichment of day 21 versus day 0 DEGs from identified lung cell subsets (FDR < 0.05). Enrichment was performed using hypergeometric distribution with BH correction. **e,** Volcano plot showing DEGs in alveolar type II epithelial cells. Significant DEGs are highlighted in red (log2FC cutoff > 0.25; FDR cutoff < 0.05). **f,** Chord diagram and heatmap showing inferred IFN-g signaling network. Edge color and arrow represents sender source and edge weights indicate interaction strength represented by computed communication probability. **g,** CD86 activation in lung innate cells after CD4 T cell or IFN-g depletion in day 21 post-BCG IV vaccinated mice. Data representative of one independent experiment with n=5-9 in CD4 depletion, and n=5 in IFN-γ depletion. **h,** Serum IFN-γ and TNF-α levels at day 19-21 in BCG IV vaccinated mice following CD4 T cell depletion. Data representative of two independent experiments (n=9-11). Statistical analysis was performed by One-way ANOVA with Tukey’s multiple comparison test; *, P < 0.05; **, P < 0.01; ***, P < 0.005; ****, P < 0.0001 (**g and h**). scRNA-seq data generated from cells pooled from n=5 mice at day 21 and 0, respectively. Abbreviations: AM, Alveolar macrophage; IM, interstitial macrophage; ATI, Alveolar type I; ATII, Alveolar type II; PNEC, Pulmonary neuroendocrine cells; SMC, smooth muscle cells; Lymphatic EC, Lymphatic endothelial cells.

Single cell transcriptomics further revealed a cluster of CD4^+^ T cells expressing mRNA encoding IFN-γ and subsets of myeloid and lung epithelial cells at day 21 expressing interferon-stimulated genes (ISGs), such as *Stat1* and *Bst2* **(Fig. 6b and Supplementary Fig. 6b)**. Upregulated differentially expressed genes (DEGs) were identified across various lung immune and non-immune subsets at day 21, suggesting a tissue-wide vaccine response **(Fig. 6c)**. We then performed overrepresentation enrichment of DEGs using blood transcriptional modules (BTMs)^34^ and Gene Ontology (GO) biological processes^35,36^. BTMs involved in viral sensing and innate antiviral immunity were enriched across myeloid cells, endothelial cells, alveolar fibroblasts, and particularly in alveolar type II pneumocytes, which are the target cells for SARS-CoV-2 and H1N1 influenza virus infection **(Fig. 6d)**. This was recapitulated in enrichment with GO terms, where modules related to cytokine-mediated signaling pathway and MHC-I processes were enriched in myeloid, endothelial, fibroblast and alveolar type II cells **(Supplementary Fig. 6c)**. Specifically, upregulated DEGs in alveolar type II pneumocytes included ISGs (*Ifi27I2a*, *Irgm2, Psmb9*), and genes involved in MHC class I complex (*H2-Q7, H2-K1*) **(Fig. 6e)**. We further validated the increase in ISG expression on a protein level using mice carrying green fluorescent protein (GFP) gene tagged to the *Mx1* allele (Mx1-GFP reporter). At day 21 post-vaccination, increased Mx1-GFP expression was significantly detected in macrophage, Ly6C^+^ monocyte and CD11b^+^ DC populations (*P* < 0.05) and in alveolar type II epithelial cells (*P* = 0.057) **(Supplementary Fig. 7a)**.

Since we have found CD4^+^ T cells and IFN-γ secretion to be critical for protective viral outcomes, we inferred potential cross-talk between CD4^+^ T cells and other cell types using CellChat, a cell-cell communication analysis tool as previously described^37^. Strikingly, type II IFN signaling from Th1 and NK cells interacted with myeloid (monocytes, macrophages, DCs), epithelial (Alveolar type I and II pneumocytes, ciliated cells), mesenchymal (alveolar fibroblast, mesothelial) and endothelial cells in the lungs **(Fig. 6f)**. We further evaluated *in vivo* expression of cytokines across various cell types in the lungs at day 23 post-vaccination using *in vivo* Brefeldin A injection, and found CD4^+^ T cells to be the key producer of IFN-γ **(Supplementary Fig. 7b)**.

To corroborate the finding that CD4^+^ T cells and IFN-γ are important in mediating the activation of innate and non-immune cells within the lungs, we measured CD86 activation on lung myeloid cells following depletion of CD4 T cells and IFN-γ with anti-CD4 and anti-IFN-γ antibodies, respectively, following BCG vaccination in C57BL/6 mice. We found a significant decrease in CD86 activation in alveolar and interstitial macrophages, Ly6C^+^ monocytes and DCs at day 21 post-vaccination following depletion of either CD4 T cells or IFN-γ **(Fig. 6g and Supplementary Fig. 8)**. In conjunction, CD4^+^ T cell depletion in vaccinated mice resulted in a significant reduction of serum IFN-γ and TNF-α levels at day 18-21 post-vaccination **(Fig. 6h)**. These results indicate that feedback from CD4^+^ T cells to immune and non-immune cells through IFN-γ signaling is essential for sustained innate activation at 3 weeks post-vaccination and could potentially be a driving factor for inducing broad protection against viruses.

### Spatial transcriptomics of CD4-dependent innate antiviral program in lungs

To further validate that CD4^+^ T cells stimulate ISG expression in lung epithelial cells and map out the spatial localization of cell populations within the lungs 21 days after vaccination, we profiled regional lung whole transcriptomes in BCG IV vaccinated mice with or without CD4^+^ T cell depletion using the GeoMx Digital Spatial Profiler (DSP) whole transcriptome atlas (WTA) assay^38^. We used fluorescently-labeled CD4 and Pan-cytokeratin (PanCK) antibodies to identify CD4 T cells and lung epithelial cells, respectively. In addition, on an adjacent serial section, we stained for BCG and myeloid cells using antibodies targeting mycobacterial antigens and CD68 to guide region of interest (ROI) selection. ROIs were selected based on the presence of CD4+ cells, PanCK+ cells, and BCG in close proximity, and a total of 29, 20, 15 ROIs were selected in BCG vaccinated, BCG vaccinated with CD4 depletion, and unvaccinated groups, respectively **(Fig. 7a, b, c)**. Notably, we detected overlapping BCG+ and CD68+ cells, indicating the presence of BCG-infected macrophages **(Fig 7b)**. As expected, a lack of CD4^+^ T cells was found in the lungs of CD4^+^ T cell-depleted mice, similarly in unvaccinated mice **(Fig. 7d, e)**. Consistently, CD4^+^ T cells detected on the GeoMx DSP displayed a high level of *Ifng* expression and genes related to T cell activation, such as *Ccl5* and *Cd48* **(Fig. 7f and g)**.

**Figure 7.**
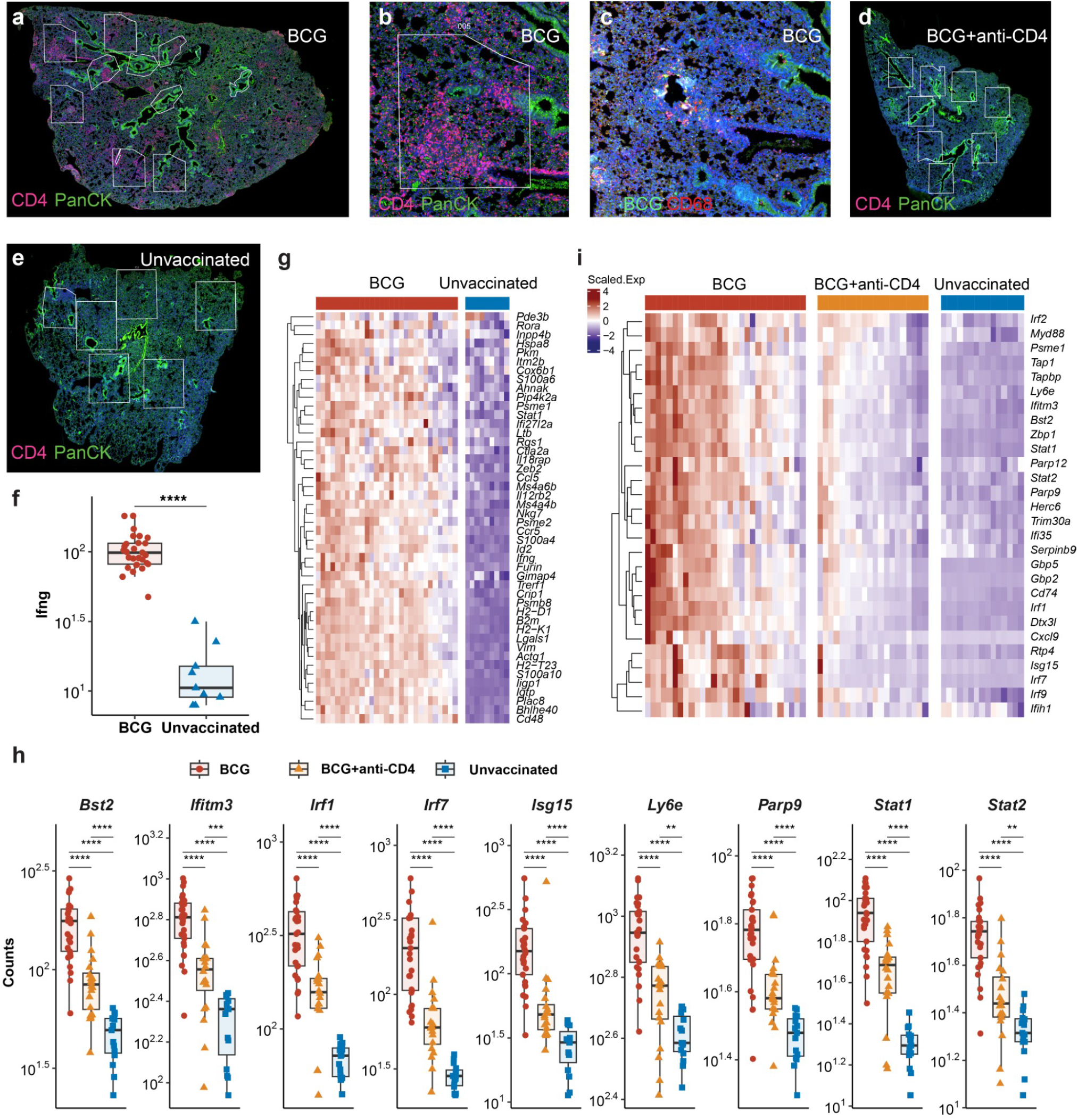
Spatial transcriptomics of CD4-dependent innate antiviral program in lungs. **a**, Representative image of GeoMx ROIs selected on BCG-vaccinated mouse lung with CD4 and pan-cytokeratin (PanCK) staining. **b,** Magnified image of representative ROI from BCG-vaccinated group. **c,** Representative image of serial lung sections stained with BCG and CD68 myeloid markers. **d and e,** Representative image of GeoMx lung sections from CD4-depleted BCG-vaccinated and unvaccinated groups, respectively. **f and g,** *Ifng* and key differentially expressed genes (DEGs) expressed in CD4+ cells in BCG-vaccinated and unvaccinated group detected from spatial transcriptomics using GeoMx. DEGs are taken from scRNA-seq data in Fig. 6. **h and i,** Interferon-stimulated gene (ISG) expression in PanCK+ epithelial cells detected from spatial transcriptomics using GeoMx. Statistical analysis was performed by Wilcoxon rank-sum test with Benjamini-Hochberg correction. **, FDR < 0.01; ****, FDR < 0.0001.

Most importantly, ROIs from the CD4^+^ T cell depleted group exhibited significantly lower expression of a variety of ISGs as compared to BCG-vaccinated control, such as *Stat1, Bst2 and Ly6e,* and these ISGs were similarly found to be upregulated in BCG-vaccinated mice in our scRNA-seq data **(Fig. 7h, 7i and 6b)**. Notably, the level of ISG expression did not reach baseline levels as found in unvaccinated mice **(Fig. 7h)**, synonymous with the presence of residual CD86 activation found in myeloid cells even after CD4^+^ and IFN-γ depletion **(Fig. 6g)**. This suggests that other IFN-γ-independent mechanisms might contribute to the reactivation of lung epithelial and myeloid cells, and led us to speculate that direct stimulation of myeloid cells by BCG, and subsequent interaction signals generated from these activated myeloid cells might be a contributing factor. Taken together, our findings strongly suggest that ISG activation within target epithelial cells, primarily driven by CD4^+^ T cell feedback, is sufficient and instrumental in eliciting non-specific viral protection.

## DISCUSSION

Our results demonstrate that intravenous vaccination with BCG confers non-specific protection against SARS-CoV-2 through a mechanism dependent on feedback of the adaptive immune response on the innate immune system, in particular antigen-specific CD4^+^ T cells imprinting an innate antiviral response in the lung. Strikingly, vaccination resulted in a biphasic pattern of innate activation characterized by an initial brief response at day 1, followed by a second prolonged response starting at between 14 and 21 days. This latter phase was synchronous with the BCG-specific Th1 response. IFN-γ secreted by such Th1 cells provided a feedback signal on the innate immune system, including myeloid cells and type II alveolar cells. Importantly, the induction of the innate and CD4^+^ T cell responses, as well as protection, was largely dependent on MyD88 signaling. Thus, these results raise the concept of “integrated organ immunity,” whereby BCG vaccination elicits a sustained antiviral program in the lung which is mediated by the dynamic interplay between the adaptive and innate immune systems, and acts in an antigen agnostic manner to provide protection against SARS-CoV-2 and influenza **(Fig. 8)**.

**Figure 8.**
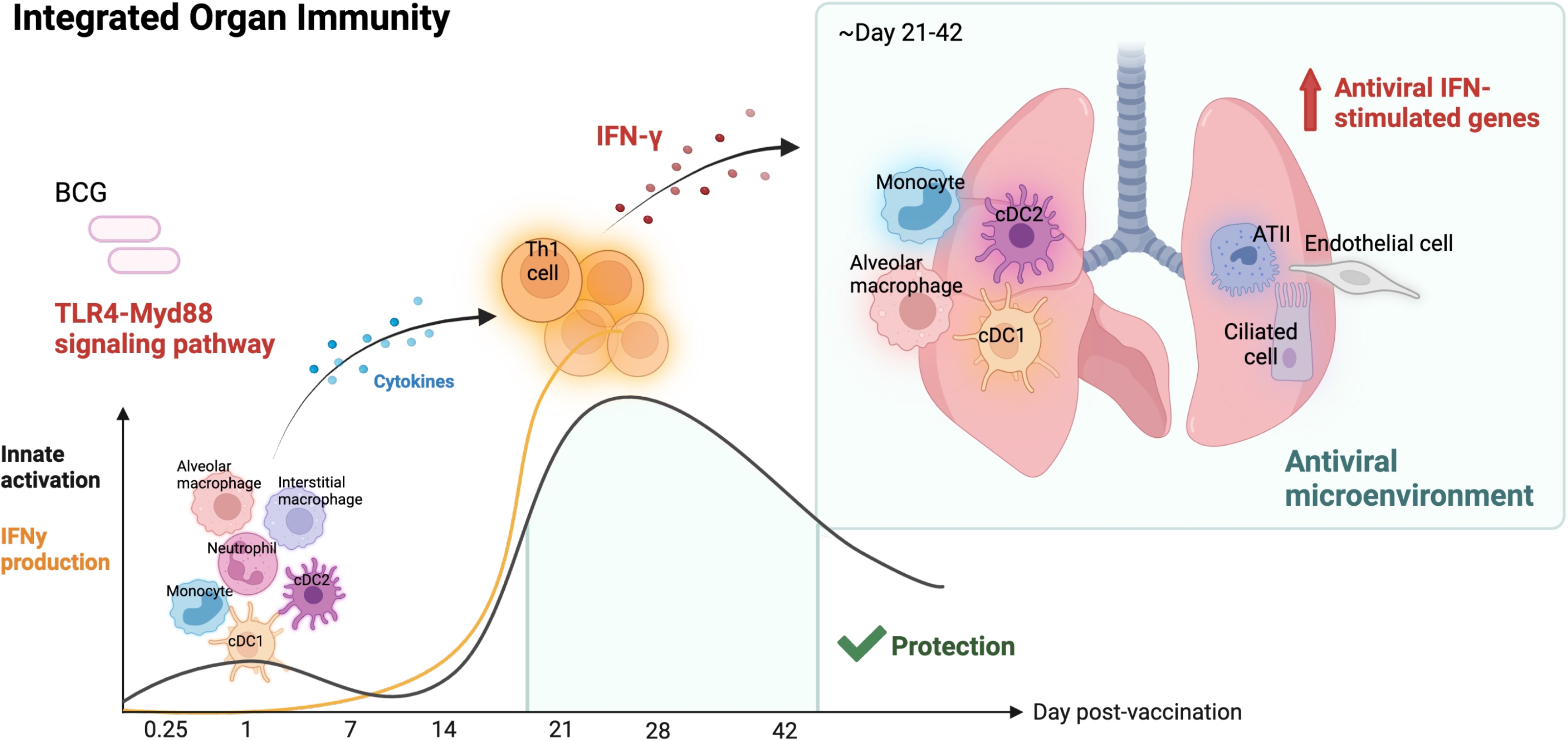
Model depicting integrated organ immunity induced by BCG, leading to non-specific protection in the lungs. BCG administered intravenously triggers the TLR4-MyD88 pathway, resulting in a biphasic pattern of systemic innate activation and production of cytokines. Second phase of sustained innate activation occurred between day 14 to 21 post-vaccination and was synchronous with upregulated BCG-specific Th1 responses at day 21. IFN-γ secreted by Th1 cells provided a feedback signal on lung myeloid and alveolar type II cells. This mechanism imprints an innate antiviral microenvironment within the lungs that is protective against SARS-CoV-2 and influenza PR8.

These results provide a shift in the prevailing paradigm that trained immunity mediated by epigenetic imprinting of innate cells such as monocytes is the chief mechanism of non-specific protection^26,27,39,40^. Several studies have reported long-lasting reprogramming or training of myeloid cells, especially with monocytes, and enhanced cytokine production associated with training^41^. In addition, our lab previously found that adjuvanted vaccines have the capacity to induce lasting antiviral gene program on an epigenetic and a transcriptomic level in myeloid cells^42,43^. Strikingly, in individuals vaccinated with an adjuvanted H5N1 vaccine several weeks previously, trained monocytes which had enhanced chromatin accessibility of loci targeted by IRFs were able to resist viral infection of dengue or zika virus following *in vitro* infection^43^. This suggests that epigenetic reprogramming of innate cells conferred non-specific protection against viruses. Additionally, Arts et al. showed in a controlled human trial that BCG vaccination led to a reduction in viremia of live yellow fever vaccine^26^. The protective outcome was primarily attributed to the enhanced secondary production of cytokines associated with trained myeloid cells^26^. Another study uncovered a T cell-independent role of BCG-induced protection against *C. albicans*, proposing that trained myeloid cells primarily contribute to the observed non-specific protection against the fungal pathogen^27^. However, in our study, we found that in the absence of CD4 T cells or IFN-γ signaling, BCG-mediated protection against SARS-CoV-2 was significantly impaired. This indicates that CD4 T cell help and IFN-γ signaling play a critical role in mediating non-specific viral protection following BCG vaccination.

In line with previous studies showing predominantly Th1 and comparatively lower Th17 responses following BCG vaccination in macaques and humans^44,45^, we found that BCG vaccination induced robust Th1 responses in the murine lungs and spleen. Furthermore, these Th1 cells are major producers of IFN-γ, which acts as a pleiotropic cytokine with immunomodulatory effects on a wide variety of cells. IFN-γ has been shown to prime various innate cell subsets, including macrophage production of anti-microbicidal products, DC antigen presentation, and NK cell cytotoxic activity^46–48^. In recent years, emerging studies have highlighted the importance of IFN-γ in maintaining long-term reprogramming or training of alveolar macrophages, with IFN-γ secretion primarily attributed to either localized CD4 or CD8 T cells^19,49^. In this context, our recent work has revealed an important role for CD8 T cell-derived circulating IFN-γ in mediating an enhanced antiviral signature in myeloid cells on day 1 following secondary vaccination with the Pfizer-BioNTech mRNA vaccine^50,51^. However, unlike what is observed in the current paper, this antiviral signature was only observed on day 1 post-boost and not afterwards, and its impact on protection against heterologous viruses is not known. Our current study revealed that myeloid cells in the lungs were reactivated 21 days following BCG vaccination and that prolonged innate activation and protection against SARS-CoV-2 was strongly dependent on feedback from CD4 T cells, in the form of IFN-γ signaling. In particular, IFN-γ signaling was found to interact with various myeloid, epithelial, and endothelial cell subsets, and notably with type II alveolar pneumocytes, which are target cells for both SARS-CoV-2 and influenza virus^52,53^. IFN-γ could exert its effects by directly inhibiting viral replication or by inducing intracellular gene programs in target cells, such as the transcription regulator IRF1 and protein kinase R (PKR), which are essential components of innate immunity against viral infections^54,55^. Collectively, our findings strengthen the role of cytokine feedback through adaptive-innate immune interactions in heterologous beneficial effects of vaccines.

BCG triggers multiple innate pathways leading to downstream immune activation. Previous studies have demonstrated the importance of MyD88 in promoting the immunotherapeutic effects of BCG against bladder cancer^56^. In addition, the IL-1R-MyD88 pathway has been implicated in the priming of skin dendritic cells (DCs) and subsequent activation of CD4 T cells in lymph nodes^57^. In our study, we demonstrated that an absence of MyD88 signaling suppressed the non-specific antiviral effects of BCG, suggesting its crucial involvement in mediating the immune responses induced by BCG vaccination. Specifically, we identified TLR4 as the key receptor responsible for eliciting MyD88-dependent immune activation following BCG vaccination. These findings suggest that vaccine strategies targeting the TLR4 pathway could potentially be harnessed in the development of vaccines aimed at providing broad protection.

The timing of immune responses following BCG vaccination appeared to be pivotal for the observed non-specific viral protection. We have shown that BCG, particularly with the BCG TICE strain used in this study, conferred a specific time window of protection against SARS-CoV-2 virus, occurring approximately 21-42 days after vaccination. Notably, this timeframe coincided with the second phase of upregulated innate responses and enhanced T cell responses in the lungs following vaccination. The distinctive biphasic kinetics of immune response following BCG vaccination is likely related to the life cycle and persistence of the BCG bacterium within the body. Previous studies have shown that BCG could persist in mice for 2-16 months, with the duration heavily influenced by the specific BCG strain used^58,59^. Given that different BCG strains could induce varied quality, magnitude, and temporal distribution of immune responses^60,61^, this could be a plausible explanation for the confounding outcomes regarding protection against SARS-CoV-2 virus observed in previous experimental studies using different BCG strains in mice^7,11^. Our findings indicate that BCG TICE specifically endows a distinct timeframe of heterologous protection in the lungs.

Furthermore, our findings indicate that the route of vaccination plays a significant role in determining the outcome of protection against heterologous viruses. Protection against Influenza A PR8 virus was achieved when BCG was administered via intravenous, intranasal, or intramuscular, but not subcutaneous routes. Similarly, protection against SARS-CoV-2 virus was afforded via intravenous or intranasal administration of BCG. This corroborates emerging studies that highlight the impact of vaccination route on the efficacy of BCG in conferring protection against mycobacterial infections or viruses. Consistently, previous studies have suggested that BCG administered through the intravenous or pulmonary mucosal routes induced more robust innate or T cell responses in the lungs, both in mice and rhesus macaques^14,19,44,62^. Interestingly, we also found that intramuscular administration of BCG resulted in a beneficial protective outcome, although not as striking as with intravenous or intranasal administration. This could potentially be explained by the deep injection of the vaccine into muscles with a rich blood supply, allowing for rapid dissemination, or could be attributed to compromised blood vessels during the vaccination process. Considering that the BCG vaccine is typically administered intradermally in humans, the significance of the spatial distribution of the vaccine in relation to its antiviral effects prompts us to reconsider the current administration method. Exploring alternative routes of BCG vaccination, such as intranasal, may hold potential for enhancing its protective efficacy against viral infections.

Finally, our study, along with others, has emphasized the importance of generating local immune responses at the site of viral challenge^19,44^. The mechanistic insights provided by our study offers a strategy for developing universal vaccines that can confer non-specific protection against diverse viruses. They suggest that mucosal delivery systems such as slow or pulsatile release kinetics nanoparticles could be used to deliver adjuvants (e.g. TLR ligands) plus an antigen to develop “universal vaccines” that exploit the dynamic interplay between the adaptive and innate immune systems to stimulate integrated organ immunity against diverse pathogens *in vivo*^63^. In this regard, it must be highlighted that since humans contain antigen-specific CD4 memory T cells against pathogens such as influenza or varicella zoster virus (VZV)^64,65^, as a result of prior infection or vaccination, such T cells could be exploited in a universal vaccine by including antigens specific for them, in order to generate Th1 cells in tissues that could imprint an antiviral program locally. It is tempting to speculate that such a mechanism, involving the recruitment of pre-existing memory T cells against VZV to orchestrate an antiviral innate program, might have been at play in the recent results showing the reduced risk of COVID-19 diagnosis and hospitalization in older adults vaccinated with the AS01b adjuvanted recombinant Shingrix vaccine^66^.

## METHODS

### Mice

C57BL/6 mice were purchased from Jackson Laboratories. All mice used were female mice aged between 8 and 14 weeks old. *Mx1^gfp^* mice were purchased from Jackson Laboratories (Strain #033219) and bred in the Stanford Research Animal Facility. *Myd88^−/−^, Tlr4^−/−^, Tlr9^−/−^, Tlr2^−/−^* mice were either obtained from colonies bred at Jackson Laboratories or purchased from Jackson Laboratories (*Myd88^−/−^,* strain #009088; *Tlr4^−/−^,* strain #029015, *Tlr9^−/−^,* strain #034449; *Tlr2^−/−^*, strain #004650). Mice used in this study were maintained under a 12h light/12h dark cycle at a temperature of ∼18–23 °C with 40– 60% humidity. All animal studies were conducted by following animal protocols reviewed and approved by the Institutional Animal Care and Use Committee of Stanford University.

### BCG vaccination

Mice were administered a single dose of 2-8×10^6^ CFU of BCG vaccine per mouse. Lyophilized BCG was first reconstituted with 1ml sterile saline and diluted accordingly. For intravenous administration, mice were injected with 100μl of BCG vaccine via tail vein injection. For intranasal administration, mice were inoculated with a total volume of 20μl of vaccine, with 10μl per nare. For intramuscular administration, mice were injected with 50μl of vaccine on the right gastrocnemius muscle. For subcutaneous administration, mice were injected with 100μl of vaccine at the base of the tail.

### Viruses and cells

VeroE6-TMPRSS2 cells were kindly provided by Barney Graham (Vaccine Research Center, NIH, Bethesda, MD). The SARS-CoV-2 B.1.351 seed stock (GISAID: EPI_ISL_890360) was provided by Andy Pekosz of John Hopkins University, Baltimore, MD. Viral stocks were grown on VeroE6-TMPRSS2 cells, and viral titers were determined by plaque assays on VeroE6-TMPRSS2 cells. VeroE6-TMPRSS2 cells were cultured in complete DMEM with puromycin at 10 mg/ml (catalog number A11138-03; Gibco). All viruses used in this study were deep sequenced and confirmed as previously described^67^.

### SARS-CoV-2 mouse infections

C57BL/6J mice were purchased from Jackson Laboratories. All mice used in these experiments were females 8 weeks of age. Stock B.1.351 virus was diluted in 0.9% Normal Saline, USP (catalog number RDI30296; MedLine). Mice were anesthetized with isoflurane and infected intranasally with virus (50 μl; 1 × 10^6^ PFU/mouse) in an animal biosafety level 3 (ABSL-3) facility. Mice were monitored daily for weight loss. All experiments adhered to the guidelines approved by the Emory University Institutional Animal Care and Use Committee.

### PR8 mouse infections

Influenza A/Puerto Rico/8/1934(H1N1) (PR8 virus) was a gift from Dr. Yueh-hsiu Chien lab. PR8 virus was propagated in Madin Darby canine kidney (MDCK) cell line (obtained from ATCC) according to protocol described in previous study^68^. PR8 virus stock was kept at −80°C until use. For intranasal inoculation, PR8 virus was diluted to 200 PFU in 20μl of PBS and 10μl was inoculated into each nare when mice were under isoflurane anesthesia.

### SARS-CoV-2 viral burden analysis

At three days post-infection, mice were euthanized with an isoflurane overdose. One lobe of lung tissue was collected in an Omni Bead Ruptor tube filled with Tripure Isolation Reagent (catalog number 11667165001; Roche). Tissue was homogenized using an Omni Bead Ruptor 24 instrument (5.15 ms, 15 s) and then centrifuged to remove debris. RNA was extracted using a Direct-zol RNA Miniprep Kit (catalog number R2051; Zymo) and then converted to cDNA using a high-capacity reverse transcriptase cDNA kit (catalog number 4368813; Thermo). To quantify RNA, IDT Prime Time gene expression master mix and were used with SARS-CoV-2 RDRP- and subgenomic-specific TaqMan gene expression primer/probe sets as previously described^69,70^. All qPCRs were performed in 384-well plates and run on a QuantStudio5 qPCR system.

### Histopathological analysis

Lungs were preserved in 10% neutral buffered formalin followed by paraffin-embedding and sectioning. Histology was performed by HistoWitz (HistoWiz, Inc). Briefly, slides were stained with H&E or subjected to IHC with an antibody specific for SARS-CoV-2 N protein (GeneTex, #GTX635686). Stained slides were read by a board-certified veterinary pathologist and scored from 0 to 5 for perivascular inflammation, bronchial/bronchiolar alveolar degeneration/necrosis, bronchial/bronchiolar inflammation, and alveolar inflammation or from 0 to 3 for extent of IHC positivity in the bronchi and the alveoli. A narrative description of leukocyte classes and histopathological findings observed was also provided.

### *In vivo* depletion with antibodies

Antibodies used for depletion are anti-CD4 (BioXcell, clone GK1.5, Cat #BE0003-1), anti-CD8β (BioXCell, clone Lyt 3.2, Cat #BE0223), rat IgG1 isotype control (BioXcell, HPRN, Cat #BE0088), rat IgG2b isotype control (BioXcell, LTF-2, Cat #BE0090), anti-mouse IFNγ (BioXcell, XMG1.2, Cat #BE0055), anti-mouse TNFα (BioXcell, XT3.11, Cat #BP0058). For T cell and cytokine depletions, mice were given 3 doses every consecutive 2 days of antibody intraperitonally, 5 days prior to either challenge or analysis. For cytokine depletion in SARS-CoV-2 challenge mice, mice were given an additional dose of anti-IFNγ or anti-TNFα 1 day post-challenge.

### Luminex assay

Blood was collected from mice at the indicated timepoints and serum was isolated by centrifugation in serum gel tubes. Luminex assay was performed on serum using Mouse 48-plex Procarta kits (EPX480-20834-901; Thermo Fisher) according to manufacturer’s protocol and modifications as previously described^51^. Each sample was measured in singlets. Plates were read on a FM3D FlexMap instrument with a lower bound of 50 beads per sample per cytokine/ chemokine.

### Flow cytometry analysis of innate cells in lungs and spleen

Lungs and whole spleen were harvested at the indicated timepoints. Briefly, lungs were prepared by enzymatic digestion with 1 mg/ml type IV collagenase and DNase I and processed using gentleMACS. Lung suspension was then passed through a 100 μm strainer and RBC lysis was performed. Spleen was harvested from mice and treated with 1 mg/ml collagenase type IV (Worthington) 20 min at 37 °C. Whole spleen tissue was then smashed with the back of a plunger against a 100 μm strainer to obtain single cell suspension. Single cells were stained with antibodies: Zombie UV^TM^ (1:200 dilution; BUV496; Biolegend #423107), anti-Ly6C (1:500; BV780; BioLegend #128041), anti-Ly6G (1:400; APC-Cy7; BioLegend #127624), anti-CD19 (1:100; BB700; BD #566411), anti-CD3 (1:100; BB700; BD #742175), anti-MHCII (1:400; AF700; eBioscience #56-5321-82), anti-CD11b (1:300; BV650; BioLegend #101239), anti-CD11c (1:400; BV421; Biolegend #117330), anti-CD86 (1:300; A647; BioLegend #105020), anti-Siglec-F (1:400; PE-CF594; BD #562757), anti-CD45 (1:200; BV610; BioLegend #103140), anti-CD169 (1:200; PE-Cy7; BioLegend #142412), anti-PDCA-1 (1:200; BUV563; BD #749275), anti-CD8a (1:200; BUV805; BD #612898), anti-CD103 (1:100; PE; eBioscience #12-1031-82), anti-NK1.1 (1:200; BV510; BioLegend #108738), anti-F4/80 (1:100; BUV737; BD #749283), Ghost Dye^TM^ Violet 510 (1:400; Tonbo Bioscience #13-0870-T100), anti-EpCAM (1:100; Biolegend 118214). Samples were then washed twice with PBS + 2% FBS + 1 mM EDTA and fixed with BD Cytofix (#554655). Samples were then analyzed on a BD FACS Symphony analyzer with BD FACS Diva v.8.0.1.

### *Ex vivo* stimulation of BCG-specific T cells

Lungs and spleen were harvested from mice at the indicated timepoints. Lungs were prepared according to a similar protocol for innate cell analysis as described above, except lungs were lung suspension was further processed with Percoll centrifugation and single cells were isolated from the interphase of a 70-40% Percoll gradient. Whole spleen was harvested and processed without enzymatic digestion. Lung or spleen single cell suspension was plated at ∼2 × 10^6^ cells per well in a 96-well U-shaped plate, and restimulated with either crude BCG at 2-8 x 10^4^ CFU or BCG purified protein derivative (PPD) at 20 μg/μl concentration.

### Intracellular cytokine staining assay for BCG-specific T cells

Lungs and spleen were harvested from mice at the indicated timepoints. Lungs were prepared by gentleMACS Lung_01_01 program, followed by incubation with collagenase IV (1mg/ml) and DNase I (Sigma) in PBS+2%FBS for 30 min. Lung suspension was then processed using gentleMACS Lung_02_01 program and passed through a 70μm strainer. Red blood cell lysis was performed with ACK lysis buffer (Quality Biological) and lung suspension was further processed with Percoll centrifugation. Single cells were isolated from the interphase of a 70-40% Percoll gradient. Whole spleen was harvested and processed without enzymatic digestion. Both lung or spleen single cell suspension was plated in complete RPMI 1640 medium at 1-2 × 10^6^ cells per well in a 96-well U-shaped plate, and restimulated with either crude BCG at 2-8 x 10^4^ CFU or BCG purified protein derivative (PPD; Cedarlane) at 20 μg/μl concentration. Cells were restimulated *ex vivo* at 37°C, 5% CO2 condition for 1.5-2h. Brefeldin-A (10 μg ml^−1^) was added and cells were incubated overnight. On day 2, cells were stained with a protocol as previously described^51^. Briefly, cells were stained with Ghost Dye Violet 510 (Tonbo Biosciences) for 10 min at RT in PBS+1mM EDTA. After washing, cells were blocked with Fc receptor antibody α-CD16/32 (clone 2.4G2, BD) for 5 min on ice before staining with fluorochrome-conjugated antibodies in FACS staining buffer (1× PBS, 2% FBS): CD3 (1:50 dilution; clone 145-2C11, BioLegend), CD8α (1:200 dilution; clone 53-6.7, BioLegend), CD4 (1:200 dilution; clone RM4-5, BioLegend), CD44 (1:400 dilution; clone IM7, BioLegend), CD69 (1:200 dilution; clone H1.2F3; BioLegend) and CD45 (1:200 dilution; clone 30-F11, BioLegend). Cells were incubated for 20 min at 4°C and washed 2x with PBS + 2% FBS + 1 mM EDTA. Cells were then permeabilized with BD Fix/Perm for 20 min at room temperature (RT) and stained intracellularly with antibodies in 1x Perm buffer (BD Biosciences): anti-IFN-γ (1:100 dilution; clone XMG1.2, BioLegend), anti-TNF-α (1:100 dilution; clone MP6-XT22, BioLegend), IL-2 (1:100 dilution; clone JES6-5H4, BioLegend) and anti-IL-4 (1:100 dilution; clone 11B11, BioLegend). Cells were then washed twice with 1x Perm buffer and fixed with BD Cytofix for 10 min at room temp. Data was acquired on BD Symphony analyzer and analyzed using FlowJo v10.

### *In vivo* staining of IFN-γ

Mice were injected with Brefeldin A (BFA; Sigma) prior to tissue harvesting as described previously^71^. Briefly, mice were injected intravenously in the tail vein with 0.5mg/ml BFA suspended in 500ul PBS. 6h after BFA injection, mice were sacrificed and lung tissues were harvested. Lungs were processed to obtain single cells according to the procedure for preparing cells for innate analysis as outlined above. Single cells were then stained with surface antibodies for innate analysis as outlined above, with additional anti-TCRg/d (1:50; PE; Biolegend #118108). Cells were then permeabilized using BD Fix/Perm for 20 min at RT and stained intracellularly with anti-IFN-γ (1:100 dilution; clone XMG1.2, Biolegend), anti-TNF-α (1:100 dilution; clone MP6-XT22, BioLegend).

### CyTOF antibody conjugation

Antibodies were conjugated to respective heavy metal isotopes with commercially available MaxPar (Fluidigm) reagents using an optimized conjugation protocol^72^. In short, antibodies were reduced with 4 mM TCEP (Thermo Fisher) for 30 min at 37°C and washed two times. For conjugations using MaxPar reagents, metal chelation was performed by adding metal solutions (final, 0.05 M) to chelating polymers and incubating for 40 min at RT. Metal-loaded polymers were washed twice using a 3-kDa MWCO microfilter (Millipore) by centrifuging for 30 min at 12,000g at RT. Antibody buffer exchange was performed by washing purified antibody through a 50-kDa MWCO microfilter (Millipore) and centrifuging for 10 min at 12,000g at RT. Partially reduced antibodies and metal-loaded polymers were incubated together for approximately 90 min at 37 °C. Conjugated antibodies were washed four times and collected by two centrifugations (2 min, 1,000g, RT) into an inverted column in a fresh 1.6-ml collection tube. Protein content was assessed by NanoDrop (Thermo Fisher) measurement; antibody stabilization buffer (Candor Bioscience) was added to a final volume of at least 50 vol/vol %, and antibodies were stored at 4 °C. Conjugations to 139La, 140Ce (Trace Sciences International) were performed following the above protocol, using Fluidigm polymers.

### CyTOF viability live-dead labeling by natural abundance cisplatin

Cisplatin (WR International, Radnor, Cat# 89150-634) was stored at −80°C as a stock solution at 100 mM in DMSO (Hybrimax, Sigma Aldrich). Working solutions (10 mM) were prepared fresh on the day of each experiment by diluting the stock solution into PBS at 4°C. Cells in suspension were centrifuged at 300×g for 5 minutes and resuspended in 1ml serum-free RPMI at 2×10^6^ cells/ml. The cisplatin working solution was added to cells at a final concentration of 25 μM for 1 min at RT, as previously described^73^. The reaction was quenched with 3ml of RPMI/10% FBS. Samples were then centrifuged at 300×g for 5 min and cell pellets were resuspended in 1 ml RPMI/10% FBS and processed for mass cytometry staining.

### CyTOF palladium barcoding and staining with heavy-metal conjugated antibodies

Palladium barcoding and staining with heavy metal-conjugated antibodies. To eliminate technical variability during staining or acquisition, individual samples within one experiment were palladium barcoded as described previously^74^ and combined into a composite sample before further processing and staining. Cell surface antibody master mix in Cell Staining Medium (CSM: low-barium PBS with 0.5 % bovine serum albumin (BSA) and 0.02 % sodium azide (all Sigma-Aldrich) was filtered through a pre-wetted 0.1-μm spin-column (Millipore) to remove antibody aggregates and added to the samples. After incubation for 30 min at RT. cells were washed once with CSM. To enable intracellular staining, cells were permeabilized by incubating with ice-cold MeOH for 10 min on ice and washed two times with CSM to remove any residual MeOH. Intracellular antibody master mix in CSM was added to the samples and incubated for 1 h at RT. Cells were washed once with CSM and resuspended in intercalation solution (1.6% PFA in PBS and 0.5 μM rhodium intercalator (Fluidigm) for 20 min at RT or overnight at 4 °C. Before acquisition, samples were washed once in CSM and twice in ddH2O and filtered through a cell strainer to obtain single cell suspension. Cells were then resuspended at 1 × 10^6^ cells per ml in ddH2O supplemented with 1× EQ Four Element Calibration.

### CyTOF analysis

Pre-processing of mass cytometry data was performed using Premessa (v0.3.4). Bead-based normalization and debarcoding were performed to obtain individual FSC files for each animal. FCS files were then imported into CellEngine (CellCarta) and single live cells gated out based on cell size and DNA content. Abundance values reported by the mass cytometer were then transformed using arcsinh with a co-factor of 5. Clustering was performed with surface markers using FlowSOM with number of clusters defined as 10^75^. Mean of abundance values was calculated for each animal and statistical test from vaccinated and unvaccinated groups was performed using Wilcoxon rank-sum test and P value < 0.05 was considered significantly different between both groups. For data visualization, column was quantile normalized for each individual marker and mean of scaled values computed and plotted for each animal using dot plot or heatmap.

### Cell isolation and library preparation for scRNA-seq

To obtain lung epithelial and immune cells, lungs were harvested and processed according to a protocol as previously described^76^. Briefly, lungs were lavaged with digestion medium containing RPMI 1640, elastase (Worthington), dispase II (Sigma) and DNase I (Sigma). Lungs were then harvested and incubated in the digestion medium for 45 min at 37°C. After incubation, lungs were cut into small pieces using scissors and incubated with RPMI 1640 containing liberase (Sigma) and DNase I for 30 min at 37°C. Dissociated cells were then passed through a 70μm strainer and RBCs lysed using ACK lysis buffer. To enrich for CD11b+, EpCAM+ and CD11c+ cells, cells were stained with surface antibodies from the innate analysis panel as outlined above, and sorted on the FACSAria II machine. Sorted cells were then combined with the non-enriched fraction at a 3:7 ratio and resuspended in cold PBS supplemented with 1% BSA (Miltenyi) and 0.5 U/μl RNase Inhibitor (Sigma Aldrich). Cells were partitioned into Gel Beads-in-emulsion (GEMs) using the 10x Chromium 3′ V3 chemistry system (10x Genomics). The released RNA was reverse transcribed in the C1000 touch PCR instrument (Bio-Rad). Barcoded cDNA was extracted from the GEMs by post-GEM RT-cleanup and amplified for 12 cycles. Amplified cDNA was subjected to 0.6x SPRI beads cleanup (Beckman, B23318). 25% of the amplified cDNA was used to perform enzymatic fragmentation, end-repair, A tailing, adapter ligation, and 10X specific sample indexing as per manufacturer’s protocol. Sequencing libraries were generated and the quality was assessed through Bioanalyzer (Agilent) analysis. Libraries were pooled and sequenced on the HiSeq 4000 instrument (Illumina) with a targeted read depth of 40,000 read pairs/cell.

### scRNA-seq analysis

scRNA-seq data obtained on the 10x genomics platform was pre-processed using Cell Ranger and raw counts were imported into R (v4.2.2) and analyzed using Seurat (v3.1.4). Briefly, raw counts from 10x genomics were filtered to remove cells greater than 10% mitochondrial reads, >50,000 and <500 RNA counts. Doublet prediction was performed using the R package DoubletFinder and predicted doublets were using removed from the dataset. Filtered read counts were scaled and transformed. The top 2000 variable RNA features were used to perform PCA on the log-transformed counts. The first 22 principal components (PCs) were selected based on a scree plot to perform clustering and UMAP projections. Clustering was performed with Seurat SNN graph construction followed by Louvain community detection with a resolution of 0.4, yielding 27 clusters. Differentially expressed genes (DEGs) at day 21 compared to baseline were identified using Seurat’s FindMarkers function and calculated by Wilcoxon rank-sum test with Benjamini Hochberg correction. Genes with an FDR < 0.05, absolute log fold change >0.25 compared to baseline and pct > 0.1 were considered significant. Gene set enrichment was performed using the clusterProfiler R package (v4.6.2). Over-representation analysis was performed with BTM gene modules and GO biological processes gene set. P-values for the overrepresentation analysis were determined by hypergeometric distribution. Pathways with FDR < 0.05 were considered significant. CellChat (v1.6.1) was used to identify ligand-receptor interaction pairs computed based on a communication probability^37^.

### GeoMx processing and analysis

Spatial profiling of mouse lung samples was performed using the NanoString Technologies GeoMx™ Digital Spatial Profiler through the NanoString Technologies Technology Access Program (TAP) in Seattle, WA, USA. The GeoMx™ Mouse Whole Transcriptome Atlas Panel was applied to paraffin-embedded sections of mouse lung tissue. 5um sections were baked for two hours at 65C for paraffin removal before loading onto a Leica BOND RX for tissue rehydration in EtOH and ddH2O, heat-induced epitope retrieval (ER2 for 20 minutes at 100°C) and proteinase K treatment (1.0 μg/ml for 15 minutes at 37°C). Tissue sections were then hybridized with the Mouse Whole Transcriptome Atlas (WTA) probes overnight at 37°C. Following 2x 5min stringent washes (1:1 4x SSC buffer & formamide), the slides with the WTA probes were blocked before morphology marker antibody staining. Unconjugated antibodies (WTA probe slides: anti-CD4 (cat#25229, Cell Signaling Technologies | serial sections: anti-BCG, AB905, AbCam) were incubated for 1 hour, before 4 2x SSC washes. Secondary antibodies (All slides: anti-rabbit with AF594, A11037, ThermoFisher) were then incubated for 30 minutes followed by 4 washes in 2xSCC. An additional blocking step, including the addition of rabbit serum, was performed before the addition of PanCK on the WTA probe slide (488 channel, ab125212, Novus Biologicals) and CD68 on the serial section (647 channel, NBP2-33200AF488, AbCam). Syto83 (532 channel, S11364, Invitrogen) was also added to slides for nuclear staining. After region of interest (ROI) selection, UV light was directed by the GeoMx at each AOI releasing the RNA-ID containing oligonucleotide tags from the WTA probes for collection into a unique well for each area of illumination (AOI). For library preparation, Illumina i5 and i7 dual indexing primers were added to the oligonucleotide tags during PCR to uniquely index each AOI. Next generation sequencing (NGS) library preparation was performed according the NanoString Technologies published protocol (GeoMx DSP NGS Readout User Manual (nanostring.com)). The prepared library was purified using AMPure XP beads (Beckman Coulter, San Jose, CA, USA), and the library quality was assessed using a Bioanalyzer (Agilent Technologies, Santa Clara, CA). The required sequencing depth was determined by calculating the sum of all of the areas of illumination (in um2) and multiplying the total area (in um2) by 100 (GeoMx DSP Library and Sequencing Guide (nanostring.com)). Sequencing was performed using a NextSeq2000. FASTQ files generated upon sequencing were converted to Digital Count Conversion (DCC) files using the NanoString Technologies publicly available pipeline (v2.0.21) (GeoMx® NGS Pipeline (illumina.com)). Through the pipeline, individual probe alignment was performed using identification of read tag sequences (RTS), and duplicate counts of probes generated during the library preparation polymerase chain reaction were removed through evaluation of the probe unique molecular identifier (UMI) sequences.

DCC files were analyzed using the publicly available NanoString Technologies GeoMx™ Digital Spatial Profiler R vignette (R package GeomxTools 3.5.0; Bioconductor v3.18; Analyzing GeoMx-NGS RNA Expression Data with GeomxTools (bioconductor.org)). The number of raw reads, the percentage of aligned reads, and the sequencing saturation of each area of illumination were evaluated to determine the quality of the library preparation. Areas of illumination were included in the analysis if they had more than 1000 raw reads, if over 80% of the reads were aligned correctly, and if the sequencing saturation was above 50%. A control sample with no target probes added, termed a “no template control,” was included in the library preparation process and sequenced to estimate library preparation contamination. Negative control probes, designed using External RNA Controls Consortium (ERCC) sequences, are included in each GeoMx™ RNA assay to estimate background noise and the limit of quantitation. For this work, the limit of quantitation was defined as the negative probe geomean plus the geometric standard deviations of the negative probes. Target probes were filtered according to their frequency in each AOI. Targets were kept in the analysis if they were detected above the limit of quantitation in >5% of areas of illumination. Targets were normalized using Q3 normalization, where individual targets were normalized to the 25% most highly counted targets of all targets. Unbiased hierarchical cluster analysis was performed using all normalized targets. Statistical analyses between groups were performed using linear mixed models corrected for random slope.

### Quantification and statistical analysis

scRNA-seq and CyTOF statistical analyses were completed as described above. Statistical analysis for other experiments was performed with Prism (GraphPad Prism v9.5.1). For comparing two groups, P values were determined using two-tailed Mann-Whitney test. For comparing multiple groups across multiple timepoints, Two-way ANOVA with Tukey multiple comparison test was performed. For comparing two groups across multiple timepoints, Two-way ANOVA with Sidak multiple comparison test was performed. For comparison of BCG vaccinated group across multiple timepoints, one-way ANOVA followed by Tukey multiple comparison test were applied. For comparison of multiple timepoints to baseline values, one-way ANOVA with Dunnett’s multiple comparison test for each timepoint compared to baseline was used. Survival analysis was performed using log-rank Mantel Cox test. Differences between groups were considered significant for P values < 0.05. Mice were assigned to the various experimental groups randomly.

## ACKNOWLEDGEMENTS

We thank the technical support from the Stanford Shared FACS Facility. Data was collected on instruments in the Shared FACS Facility purchased by Parker Institute for Cancer Immunotherapy (PICI) or obtained using NIH S10 Shared Instrument Grant (Symphony: 1S10OD026831-01). We thank the Stanford Functional Genomics Facility for technical assistance; Dhananjay Wagh and Ed Kim for library preparation, John Coller for data analysis. The sequencing data was generated with instrumentation purchased with NIH funds (S10OD025212 and 1S10OD021763) at the Stanford Genomics Facility. We thank the Stanford Human Immune Monitoring Core (HIMC) for technical assistance in performing Luminex assays. We thank Benjamin Franco from Stanford Veterinary Service Center (VSC) for technical support. Diagrams were created with BioRender.com.

## AUTHOR CONTRIBUTIONS

A.L. and B.P. designed the study, interpreted the data and wrote the manuscript. A.L. led and planned the research and analyzed the data. K.F. and M.S. performed all SARS-CoV-2 mouse experiments including challenge and viral load quantification. S.W. performed mouse injections and helped with tissue harvesting and processing. Z.F. contributed to scRNA-seq analysis. T.Z.K. helped with CyTOF staining. D.S., A.D.R., Y.L. and A.P. contributed to and performed GeoMx. H.H. and V.L. helped with tissue harvesting and processing. C.L. helped with propagating PR8 virus. P.A. provided guidance and advice. G.P.N. provided CyTOF instrument.

