## Supplementary Figures for "Integrated Organ Immunity: Antigen-specific CD4-T cell-derived IFN-γ induced by BCG imprints prolonged lung innate resistance against respiratory viruses"

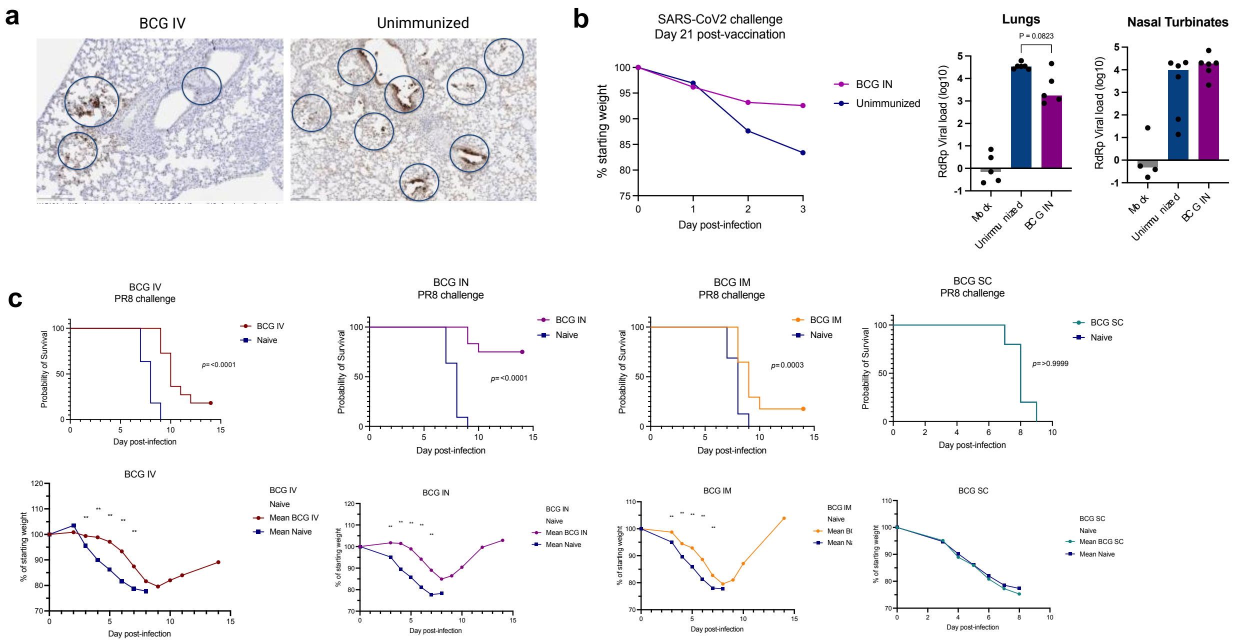


**Supplementary Fig 1. BCG delivered via multiple routes of vaccination conferred protection against SARS-CoV-2 and influenza A PR8. a,** Representative immunohistochemisry images depicting staining of SARS-CoV-2 nucleocapsid protein (NP). Blue circles represent expression of SARS-CoV-2 in focal alveoli, alveolar macrophages, alveolar pneumocytes and rare bronchiolar epithelial cells. BCG IV: Intravenous BCG vaccinated. **b,** Left, weight loss of intranasal BCG vaccinated mice followed 3 days post-SARS-CoV-2 challenge. Right, SARS-CoV-2 RNA-dependent RNA polymerse (RdRp) viral load as fold-change over mock-infected, in the lungs and nasal turbinates. Statistical analysis was performed by Mann-Whitney test. **c,** Survival plot and weight loss of mice following PR8 infection. Mice were vaccinated via various routes including intravenous (IV), intranasal (IN), intramuscular (IM), subcutaneous (SC). BCG IV and IN for PR8 infection, data combined from 2 independent experiments (n=11-12); BCG IM for PR8 infection, data combined from 3 independent experiments (n=16-17). BCG SC, data from one independent experiment. Survival analysis was performed using log-rank (Mantel-Cox) test. Statistical analysis for weight loss was performed by Two-way ANOVA with Sidak’s multiple comparisons test. **, P < 0.01.

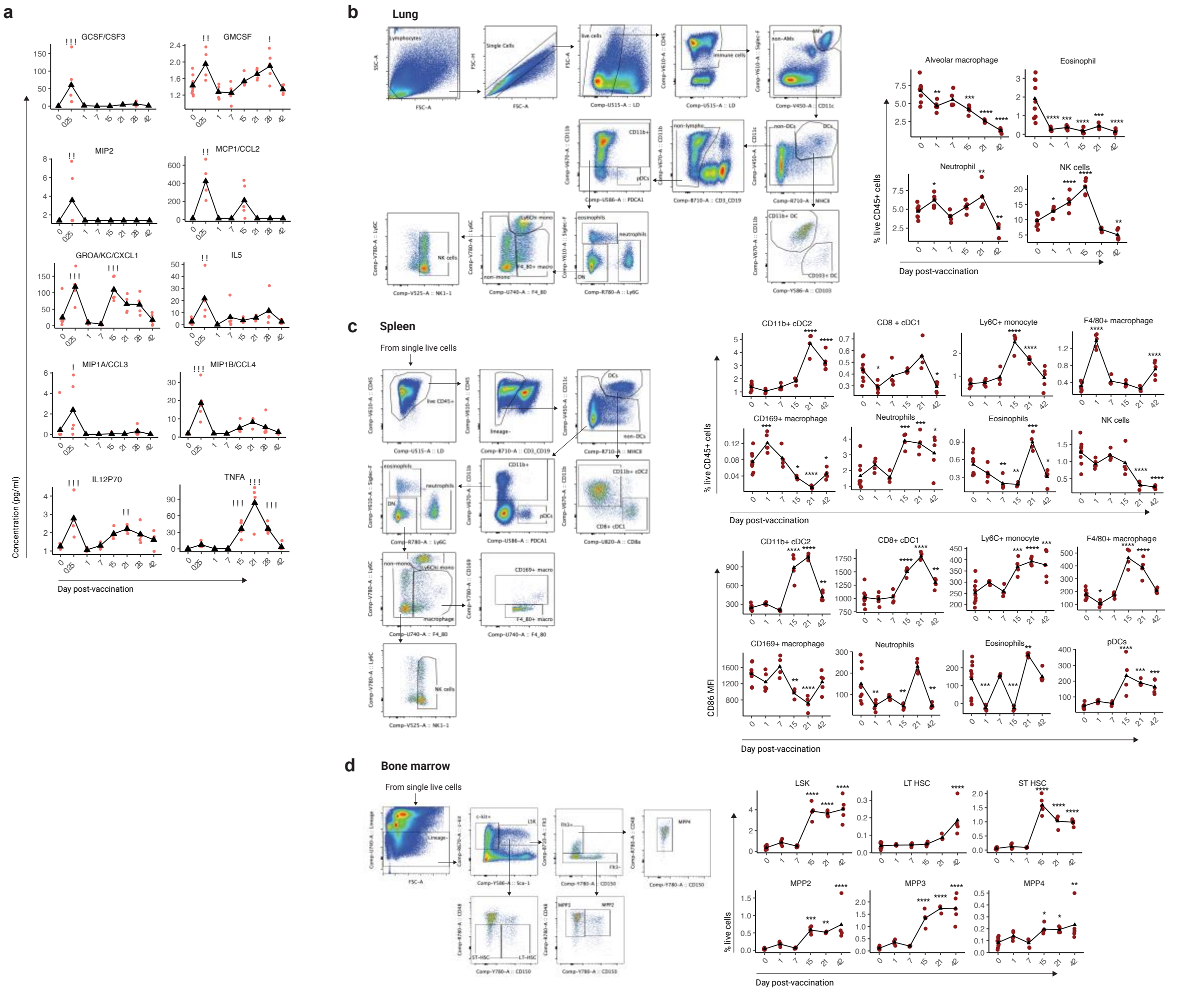


**Supplementary Fig 2. BCG induced prolonged and systemic innate activation and bone marrow stem and progenitor expansion. a,** Extended data of kinetics of serum cytokine responses measured by Luminex. **b and c,** Cell frequencies (CD45+ live cells) and activation marker (CD86 MFI) on innate cells in lungs **(b)** and spleen **(c)**, as measured by flow cytometry staining. Data is representative of two independent experiments for day 21 and one representative experiment for other timepoints (n=5-8). **d,** Frequencies of bone marrow cells (% of live cells). Data from one representative experiment. Abbreviations, LSK: Lin$^{-}$Sca-1$^{+}$c-Kit$^{+}$ cells, LT-HSC: Long-term HSC. ST-HSC: Short-term HSC. MPP: Multipotent progenitor. One-way ANOVA with Dunnett's multiple comparison test for each timepoint compared to day 0 **(b).** *, P < 0.05; **, P < 0.01; ***, P < 0.005; ****, P < 0.0001.

| Antibody target | Metal Isotope | Clone | Vendor |
|---|---|---|---|
| Phospho-IRF-3(S396) | 147Sm | 4D4G | Cell Signaling Tech |
| iNOS | 164Dy | CXNFT | ThermoFisher |
| NF-kB p65 (Ser536) | 171Yb | 93H1 | Cell Signaling Tech |
| Phospho-IRAK4 (T345/S346) | 173Yb | D6D7 | Cell Signaling Tech |
| p-JAK2 (Y1007/Y1008) | 141Pr | C80C3 | Cell Signaling Tech |
| pSTAT1 (Y701) | 143Nd | 4a | BD Biosciences |
| p-p38-MAPK (T180/Y182) | 149Sm | 36/p38 | BD Biosciences |
| pSTAT3 (Y705) | 151Eu | 4/P-STAT3 | BD Biosciences |
| p44/42 MAPK/p-ERK1/2 (T202, Y204) | 152Sm | 20A | BD Biosciences |
| pSTAT6 (Y693) | 154Sm | J71-773.58.11 | BD Biosciences |
| pSTAT4 (Y693) | 155Gd | 38/p-Stat4 | BD Biosciences |
| p-CREB (S133) | 161Dy | 87G3 | Cell Signaling Tech |
| pSTAT5 (Y694) | 165Ho | 47/Stat5(pY694) | BD Biosciences |
| IkBa amino-terminal | 166Er | L35A5 | Cell Signaling Tech |
| Phospho-TBK1/NAK (Ser172) | 168Er | D52C2 | Cell Signaling Tech |
| p-AKT (S473) | 169Tm | D9E | Cell Signaling Tech |
| CD103 | 113Ln | 2E6 | BioLegend |
| CD117 (c-kit) | 115Ln | 2B8 | Fluidigm |
| CD24 | 139La | M1/69 | BioLegend |
| CD135 (Flt3) | 142Nd | A2F10 | ThermoFisher Scientific |
| F4/80 | 144Nd | BM8 | Fluidigm |
| CD3 | 145Nd | 17A2 | BioLegend |
| CD169 | 146Nd | 3D6.112 | BioLegend |
| CD11b | 148Nd | M1/70 | Fluidigm |
| CD4 | 150Nd | RM4-5 | BioLegend |
| PDCA1 | 153Eu | 927.0 | BioLegend |
| CD11c | 156Gd | N418 | BioLegend |
| CD19 | 157Gd | 6D5 | BioLegend |
| SiglecH | 158Gd | 551.0 | BioLegend |
| CCR2/CD192 | 159Tb | 475301.0 | R&D Systems |
| CX3CR1 | 160Gd | SA01F11 | Biolegend |
| Ly6C | 162Dy | HK1.4 | Fluidigm |
| XCR1 | 163Dy | ZET | BioLegend |
| CD8 | 167Er | 53-6.7 | BioLegend |
| NK1.1 | 170Er | PK136 | Fluidigm |
| Ly6G | 172Yb | 1A8 | Fluidigm |
| Siglec-F | 174Yb | E50-2440 | BD Pharmingen |
| CD86 | 175Lu | GL-1 | Biolegend |
| CD45 | 89Y | 30-F11 | Fluidigm |
| CD64 | 140Ce | X54-5/7.1 | Fluidigm |
| MHCII | 176Yb | M5/114.15.2 | Biolegend |

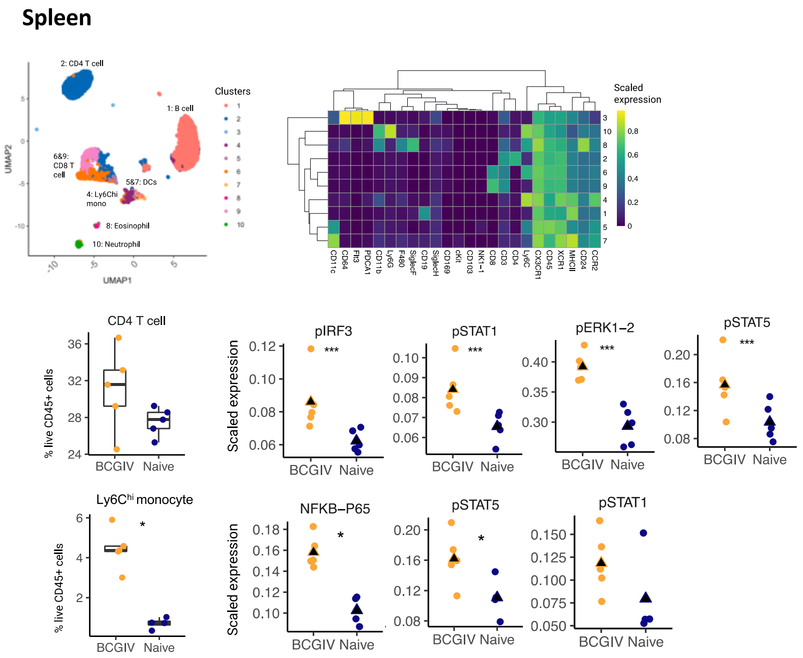


**Supplementary Fig 3. Extended data on CyTOF analysis in the spleen.** (Left) CyTOF panel used in both lung and spleen staining. (Right) UMAP embedding of the cell clusters identified by FlowSOM clustering, heatmap depicting the normalized values of key surface markers across identified clusters. (Bottom) Frequencies of CD4 T cell and Ly6Chi monocyte in the spleen and normalized values of intracellular markers detected by CyTOF. Statistical analysis was performed by Wilcoxon rank-sum test. *, $P < 0.05$; ***, $P < 0.005$.

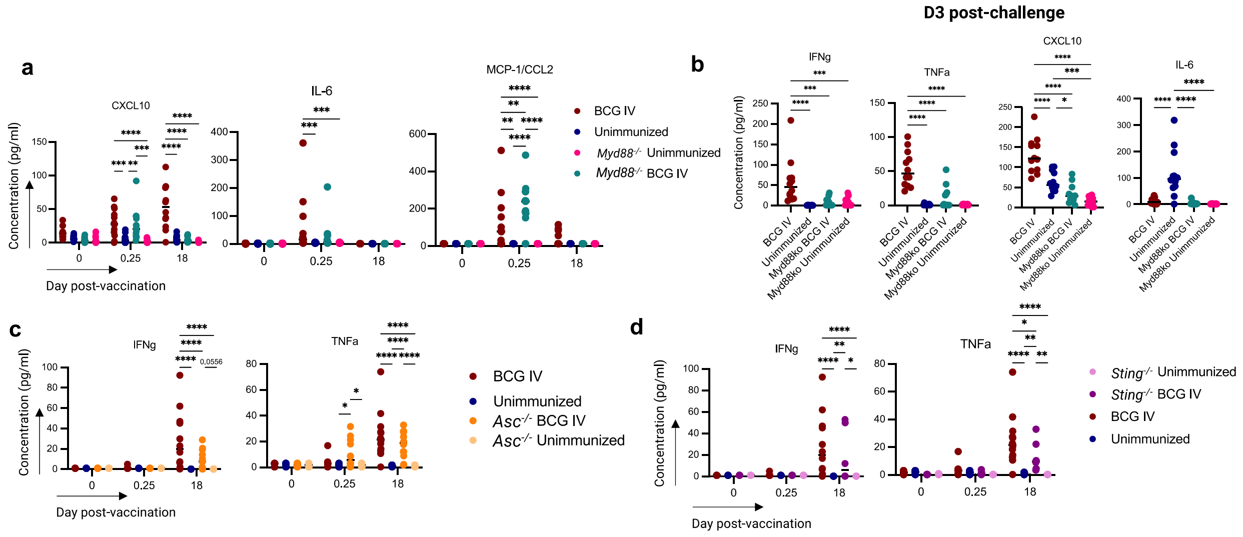


**Supplementary Fig 4. Differential immune responses to BCG vaccination in knock-out (KO) mice. a and b,** Serum cytokine responses in wild-type (WT) and *Myd88*$^{-/-}$ mice vaccinated with BCG intravenously at baseline, 6h and 18 days post-vaccination **(a)**, and day 3 post-SARS-CoV-2 challenge **(b)**. **C and d,** Serum cytokine responses in wild-type (WT) and *ASC*$^{-/-}$ **(c)** and *STING*$^{-/-}$ **(d)** mice vaccinated with BCG intravenously. Data representative of two independent experiments for *Myd88*$^{-/-}$ and *ASC*$^{-/-}$ (n=6). Two-way ANOVA with Tukey's multiple comparison test (panel b, e, f). Multiple Mann-Whitney with Holm-Sidak multiple comparisons test (panel c, d). *, P < 0.05; **, P < 0.01; ****, P < 0.0001.

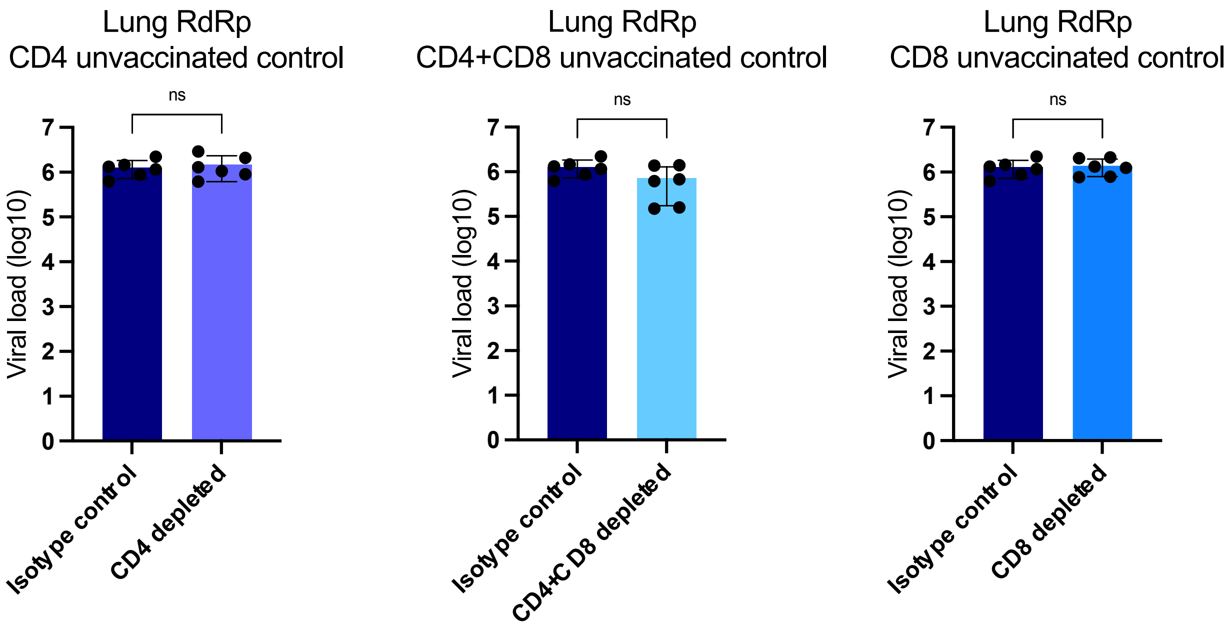


**Supplementary Fig 5. SARS-CoV-2 RNA-dependent RNA polymerase (RdRp) level in CD4 and/or CD8 antibody depleted mice.** Viral load was measured at day 3 post-challenge by quantitative PCR and is represented as fold-change over mock–infected mice (n=5-6). Statistical analysis by Mann-Whitney test; ns, not significant.

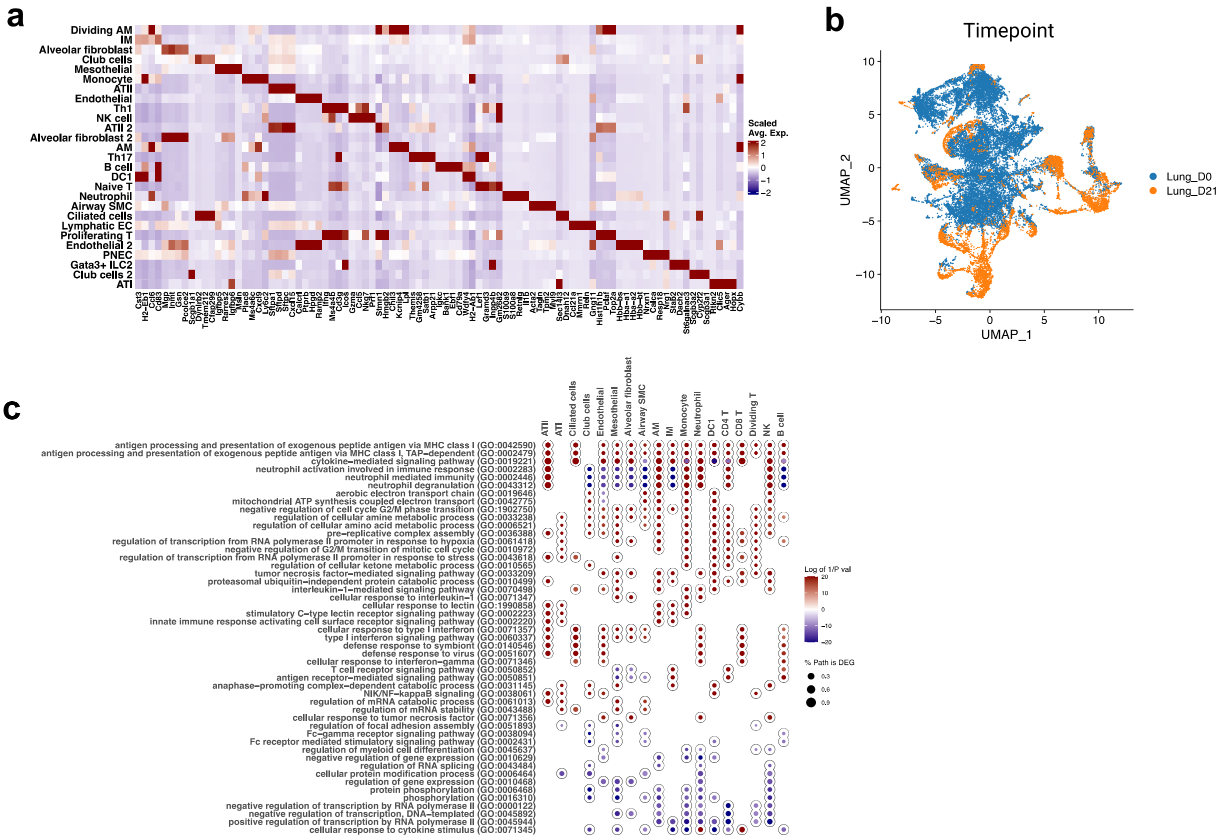


**Supplementary Fig 6. Extended data from scRNA-seq analysis of lungs in mice at day 0 and 21 post-vaccination. a,** Top variable genes defining cell clusters identified from scRNA-seq data. **b,** Distribution of cells identified day 0 and 21 post-vaccination on the UMAP embedding. **c,** Overrepresentation analysis and enrichment of Gene Ontology (GO) biological processes across identified cell clusters, using differentially expressed genes (DEGs; log2FC cutoff > 0.25; FDR cutoff  < 0.05).

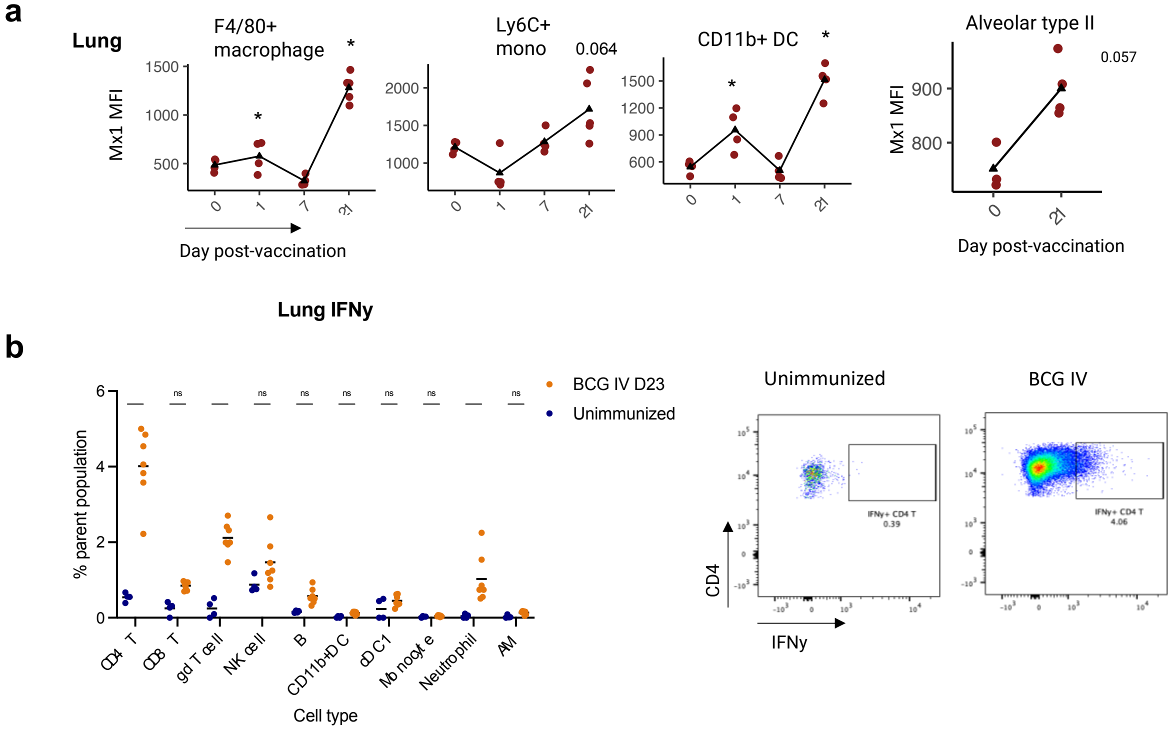


**Supplementary Fig 7. Protein level expression of interferon-stimulated gene (Mx1) and IFN-γ in lung cells, and CD4 T cells, respectively. a,** Mx1 median fluorescent intensity (MFI) of lung myeloid and alveolar type II epithelial cells detected using Mx1-GFP reporter mice at various timepoints post-vaccination with BCG intravenously (n=3-5). Data representative of two independent experiments in lung myeloid cells and one representative experiment in alveolar type II cells. **b,** Frequency of IFN-γ+ cells (% of cell population of interest) in vivo, as measured by flow cytometry staining 6h following Brefeldin A injection in mice. Data from one representative experiment (n=7). Statistical analysis was performed by Two-way ANOVA with Dunnet's test **(a;** myeloid cells**)** Sidak's multiple comparisons test **(b)**, and with Mann-Whitney test (**a**; alveolar type II cells). *, P < 0.01; ****, P < 0.0001.

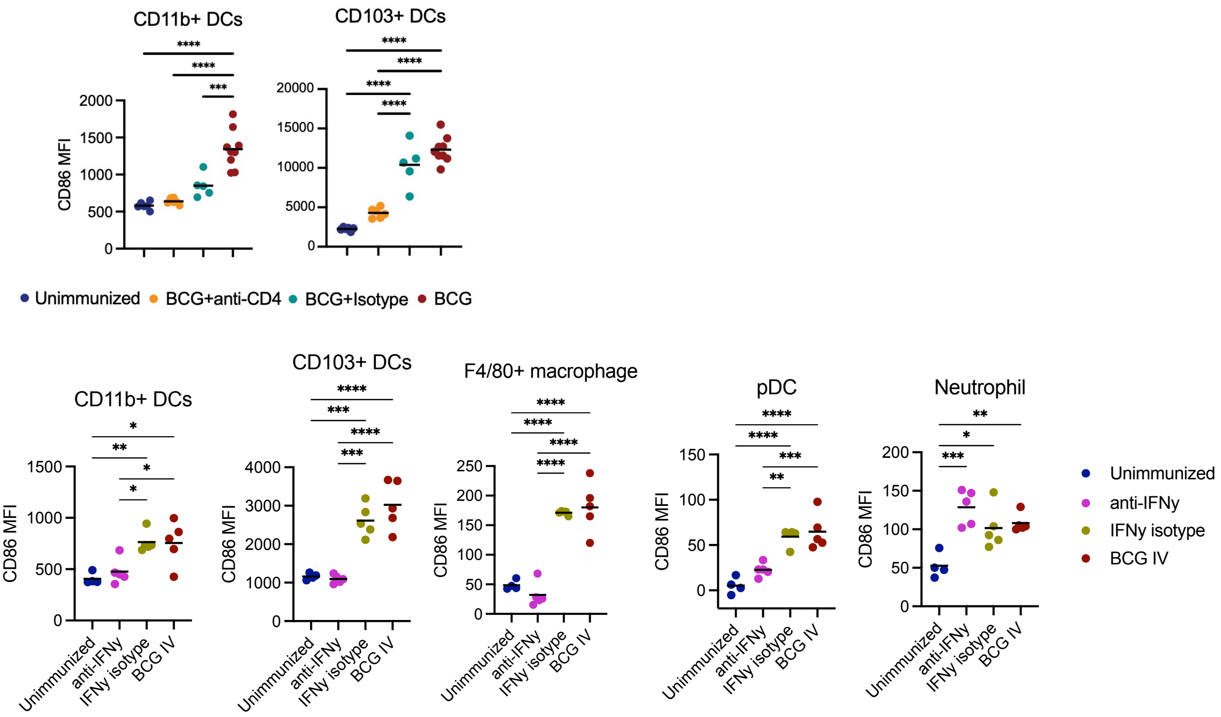


**Supplementary Fig 8. Extended data showing impaired activation of lung myeloid cells following CD4 T cell or IFN-γ depletion.** CD86 median fluorescence intensity (MFI) of DCs in the lungs at day 21 post-BCG IV vaccination following CD4 depletion (top) and IFN-γ depletion (bottom). Data representative of one independent experiment with n=5-9 in CD4 depletion, and n=5 in IFN-γ depletion. Statistical analysis was performed by One-way ANOVA with Tukey's multiple comparison test. *, $P < 0.05$; **, $P < 0.01$; ***, $P < 0.005$; ****, $P < 0.0001$.
